# Efficient small-scale conjugation of DNA to primary antibodies for multiplexed cellular targeting

**DOI:** 10.1101/662791

**Authors:** Glenn A.O. Cremers, Bas J.H.M. Rosier, Roger Riera Brillas, Lorenzo Albertazzi, Tom F.A. de Greef

## Abstract

The combination of the specificity of antibodies and the programmability of DNA nanotechnology has provided the scientific community with a powerful tool to label and unambiguously distinguish a large number of subcellular targets using fluorescence-based read-out methods. While primary antibodies are commercially available for a large class of targets, a general stoichiometric site-specific DNA labeling strategy for this affinity reagent is lacking. Here, we present a universal, site-selective, conjugation method using a small photocrosslinkable protein G adaptor that allows labeling of antibodies of different host species with a controlled number of short oligonucleotides (ODNs). Importantly, we illustrate that this conjugation method can be directly performed on commercially-available primary antibodies, on a small scale and without cross-reactivity towards other proteins, such as bovine serum albumin. In addition, we present a general, benchtop-compatible strategy to purify DNA-labeled antibodies without loss of function. The application of protein G-ODN labeled primary antibodies is demonstrated by employing three well-known methods for detecting subcellular targets using fluorescent read-out, including flow cytometry, DNA-PAINT, and dSTORM. This work thus establishes a general and efficient platform for the synthesis of a library of unique ODN-antibody conjugates, facilitating the broader use of DNA-based programmable tags for multiplexed labeling to identify subcellular features with nanometer-precision, improving our understanding of cellular structure and function.

## Introduction

In order to unravel the structure, organization, and function of subcellular components in a crowded environment, specific orthogonal labeling of a large variety of biomolecules inside the cell is essential. Currently, antibodies are the preferred affinity reagents for visualization of subcellular components, since they offer exquisite control over specificity and are commercially available for a large class of targets. The predictability of DNA nanotechnology has provided a powerful tool for providing antibodies with unique, programmable labels that allow detection via various fluorescence-based read-out methods.^1^ In general, read-out methods based on DNA have the advantage that their coding capacity, in addition to a number of spectrally distinct fluorescent tags, relies on the complementarity of unique oligonucleotide (ODN) sequences, increasing the number of labels that can be used simultaneously.^2,3^ As a result, multiple fluorescent based read-out methods are available which rely on reversible binding of short imager strands^4,5^, affinity-mediated signal amplification^6,7^, and DNA strand displacement.^8–10^ In addition, advances in the field of DNA nanotechnology have provided a powerful tool for the design of well-defined nanostructures, the DNA origami technique.^11–13^ This development has allowed the design of nanostructures which facilitate control over optical properties (e.g. brightness, color) by site-specific incorporation of fluorescently labeled ODNs.^14,15^ Combining the programmability of DNA nanotechnology with the specificity of antibody labeling therefore facilitates the design of programmable fluorescent tags that have the ability to label >100 subcellular targets and can be distinguished unambiguously.

Although primary antibodies are widely commercially available, a general stoichiometric site-specific ODN labeling strategy is missing. Traditionally, antibodies are functionalized with a modified ODN that targets chemical groups present in the native antibody (e.g. thiols and primary amines).^16,17^ However, this method lacks site-specific and stoichiometric control and can therefore result in antibodies with diminished binding capacity.^18^ Moreover, commercially available antibody solutions contain protein stabilizers, bovine serum albumin (BSA) in particular, which carry numerous functional groups that directly compete for reaction with the functionalized ODN. Several methods have been introduced to address these limitations, involving the introduction of non-canonical amino acids^19^ or specific labeling tags, including Snap-tags^20^, HaloTags^21^ and CLIP-tags.^22^ Additionally, coupling methods targeting specific regions on the antibody have been applied.^23,24^ However, these methods require genetic re-engineering of the antibody, are limited by the specific host species or sub-type of the antibody or are performed in the absence of stabilizing proteins.

Here we present a general, benchtop-compatible strategy to site-specifically label and purify commercially available primary antibodies with short oligonucleotides. Importantly, we confirm that this labeling method is selective for antibodies and shows no cross-reactivity towards other proteins. This selectivity is achieved using an ODN-functionalized protein G adaptor^25^ (pG-ODN) that can be photo-cross-linked specifically to the fragment crystallizable (Fc) region of a native immunoglobulin G-type (IgG) antibody (Fig. 1). We previously developed, and successfully used, this strategy to decorate DNA nanostructures with antibodies and Fc functionalized proteins.^26^ In this study, we optimized the ODN coupling efficiency to protein G which allowed direct conjugation of unpurified pG-ODN constructs to a native antibody, making multiplexed antibody labeling efficient and less time-consuming. We show that this strategy is compatible with antibodies from 3 different host species, including mouse IgG2a and Rabbit IgG, which together cover ≥80% of the commercially available primary antibodies.^27^ In combination with a universal, benchtop-compatible, purification method we report on the successful labeling and purification of a subset of primary antibodies. To illustrate the universal applicability of the coupling strategy, cellular labeling with pG-ODN-antibody constructs is evaluated using flow cytometry and super-resolution microscopy techniques, including stochastic optical reconstruction microscopy (STORM)^28^ and DNA point accumulation for imaging in nanoscale topography (DNA-PAINT).^2^ Our results show successful cellular labeling using the pG-ODN-antibody conjugates and underline that covalent coupling of the pG-ODN construct to the antibody is required when multiplexed cellular labeling is performed.

**Figure 1.**
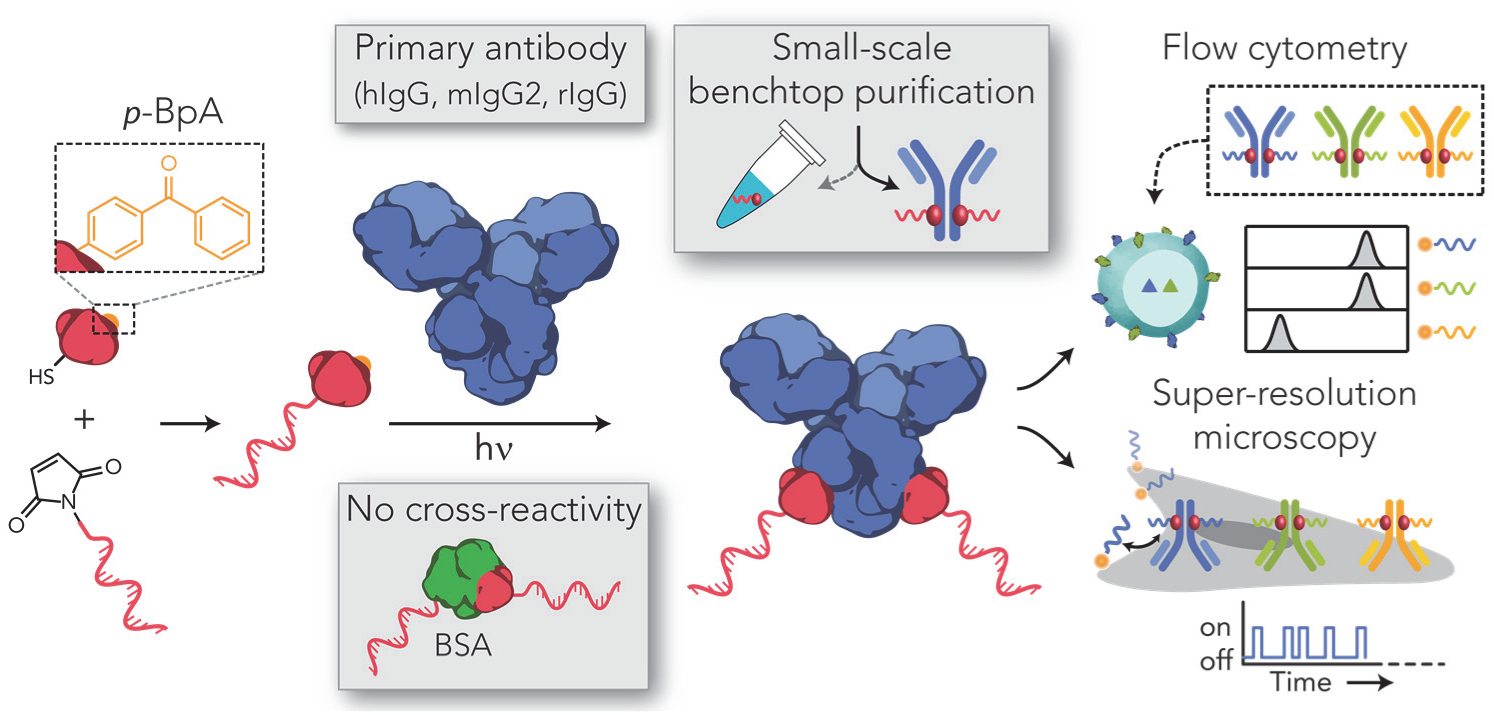
Schematic overview of the labeling strategy for site-specific functionalization of primary antibodies with an oligonucleotide (ODN) via a small protein G adaptor without cross-reactivity towards stabilizing proteins (e.g. bovine serum albumin, BSA). Protein G is functionalized and expressed with a cysteine coupled to a maleimide-functionalized ODN. Additionally, the protein G variant contains a non-natural amino acid, p-benzoylphenylalanine (p-Bpa), that covalently couples the protein G-ODN conjugate to the fragment crystallizable (Fc) region of a primary antibody, including human IgG (hIgG), mouse IgG2 (mIgG2) and rabbit IgG (rIgG), using long-wavelength UV light (365 nm). Antibody-ODN conjugates are purified via a benchtop-compatible method and can be directly used for multiplexed cellular labeling using either flow cytometry or super resolution microscopy.

## Results and discussion

To facilitate the efficient synthesis of multiple unique pG-ODN constructs, we first focused on a labeling method to couple maleimide-functionalized ODNs to protein G with a high yield. Previously, it was found that the N-terminal cysteine formed an unreactive thiazolidine adduct during pG expression, which limited the pG-ODN conjugation efficiency to ∼15%.^26^ We hypothesized that the introduction of an additional amino acid before the N-terminal cysteine would resolve this problem and increase the conjugation efficiency. Indeed, the site-specific insertion of a serine N-terminal to the cysteine showed no adduct formation after pG expression (Fig. S1). As a result, the pG-ODN coupling efficiency was increased to >90% when pG was incubated with 5-fold molar excess of maleimide-functionalized ODN (Fig. S2). Subsequently, we investigated if the pG-ODN reaction mixture could directly be used for antibody coupling without initial purification of the pG-ODN construct. To this end we incubated Cetuximab, a monoclonal IgG1 antibody, with different molar equivalents of pG-ODN. We compared the formation of pG-ODN-antibody conjugates using the unpurified pG-ODN reaction mixture and purified pG-ODN. SDS-PAGE analysis was used to monitor product formation and gel band intensity analysis showed that >80% of the heavy chains was successfully labeled with pG-ODN using a 10-fold molar excess of unpurified pG-ODN (Fig. 2a). In comparison, we observe >90% labeling efficiency when purified pG-ODN was used (Fig. S3). Additionally, lyophilization of the pG-ODN constructs did not decrease the reported conjugation efficiency, making these constructs ideal for long-term storage and shipping (Fig. S4). We note that the antibody conjugation efficiency (∼80%) does not match the yield observed after pG-ODN coupling (>90%), however, we hypothesize that pG-antibody binding is preferred over pG-ODN-antibody coupling due to the steric hindrance and electrostatic repulsion induced by the ODN. Nevertheless, based on the observed Fc labeling efficiency for unpurified pG-ODN we expect that >95% of all the antibodies (2 Fc domains) contain at least 1 ODN sequence.

**Figure 2.**
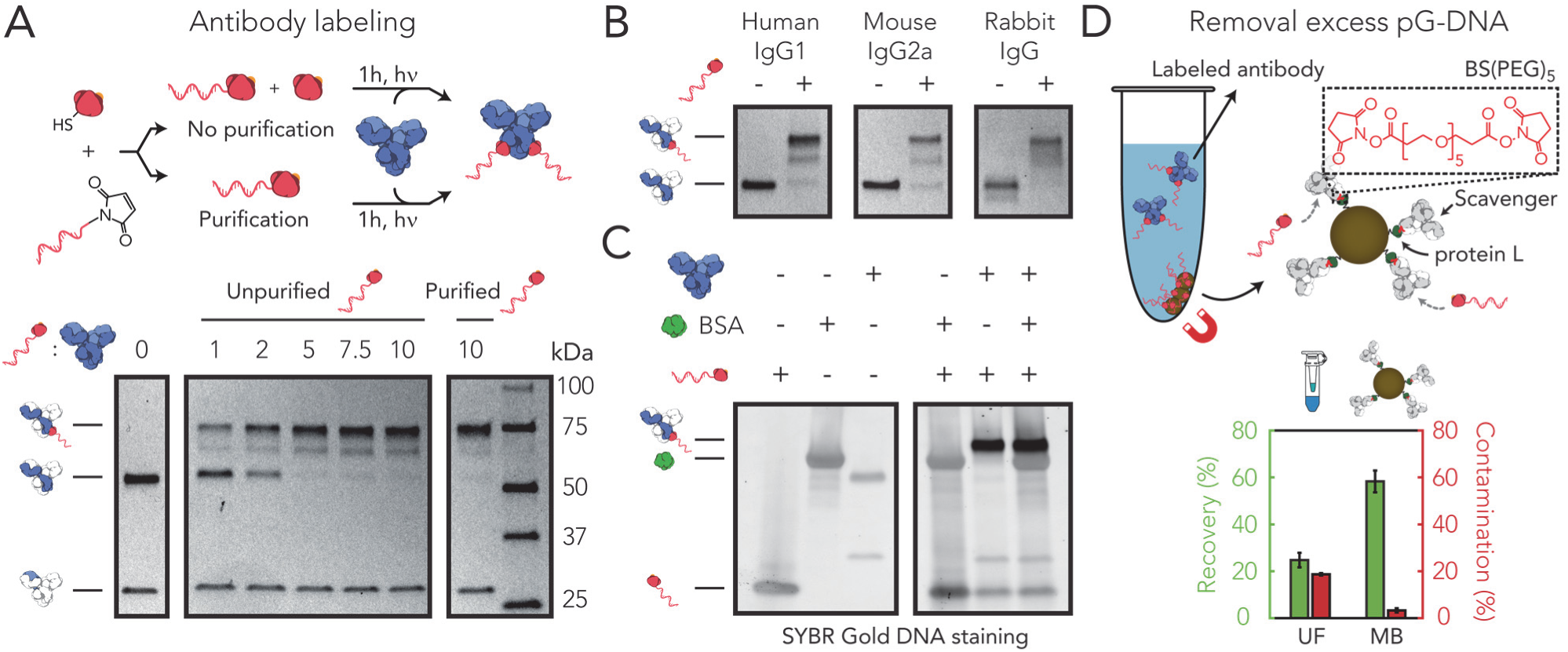
Antibody-ODN labeling and purification. (A) Protein G (pG) was conjugated to a 20 nt 3’-maleimide-functionalized ODN. Human IgG1 (hIgG1) was incubated with varying molar equivalents of unpurified pG-ODN and the labeling efficiency was analyzed and compared to purified pG-ODN under reducing SDS-PAGE conditions. (B) Reducing SDS-PAGE analysis of various IgG subclasses coupled to 10 equivalents pG-ODN and a (C) hIgG1 antibody coupled to pG-ODN in the presence and absence of 0.07% (w/v) bovine serum albumin (BSA). (D) Excess of pG-ODN was removed using protein L functionalized magnetic beads covalently coupled to a scavenger antibody via a PEGylated bis(sulfosuccinimiyl)suberate linker (BS(PEG)_5_). The average recovery (green) and contamination (red) of the magnetic beads (MB) were compared to ultrafiltration (UF). Error bars represent SD (n=3).

To examine the general applicability of the pG-ODN labeling strategy, we evaluated the coupling efficiency for antibodies of different host species. Previously, it was shown that pG was also able to photo-cross-link to mouse IgG2a and Rabbit IgG.^25^ Rabbit IgGs in particular are an important class of antibodies, since rabbits are the host species in which most primary antibodies are raised.^27^ Incubating both mouse IgG2a (mIgG2a) and rabbit IgG with 10-fold excess of pG-ODN showed successful antibody coupling, emphasizing the universal applicability of pG-ODN labeling (Fig. 2b).

Since commercially available primary antibodies are typically supplemented with protein stabilizers (e.g. bovine serum albumin (BSA), glycerol, sodium azide), pG-ODN antibody labeling should be compatible with the presence of these additives. BSA, in particular, is known to interfere with more classic ODN-labeling methods targeting functional groups in the antibody (e.g. lysine, cysteine). Commercially available kits can be used to remove BSA, however these protocols rely on extensive washing steps, resulting in extensive loss of antibody. The exclusive specificity of pG-ODN towards the Fc region should overcome this problem, making the need for BSA-free antibody solutions redundant. To test this, we performed the coupling of an antibody to pG-ODN in the presence of 0.07% (w/v) BSA, corresponding to 1% (w/v) BSA in an antibody solution of 1 mg/mL. As expected, labeling of Cetuximab in the presence of BSA did not result in cross-reactivity, confirming the specificity of pG-ODN constructs (Fig. 2c). Additionally, we evaluated the coupling efficiency of pG-ODN in the presence of other common antibody additives, including TWEEN-20, glycerol and sodium azide. Sodium azide is known to quench the reactive triplet state of *p*-Bpa, and is therefore likely to inhibit the pG-ODN crosslinking.^29^ While TWEEN-20 did not alter the coupling efficiency, both glycerol and sodium azide inhibited the formation of pG-ODN-antibody constructs (Fig. S5). However, buffer exchange of the antibody using ultrafiltration to a glycerol/sodium azide free buffer did restore the coupling efficiency. From this, we conclude that the pG-ODN labeling strategy can be used in combination with commercially available antibodies.

We next focused on the development of a universal purification approach to remove excess pG-ODN. To this extent we systematically evaluated different purification methods which either focus on size-based separation or are specifically designed for antibody purification. First, we tested the separation of pG-ODN-antibody constructs and pG-ODN using size-exclusion chromatography (SEC). The relatively large difference in molecular weight between pG-ODN and labeled antibodies, 15 and 180 kDa respectively, should provide enough separation to effectively purify the pG-ODN-antibody. To this end we applied a high-performance liquid chromatography (HPLC) system with a size exclusion column that has a fractionation range between 5 x 10^3^ and 1.25 x 10^6^ Da. Using this approach, the pG-ODN-antibody conjugate was well separated from unreacted pG-ODN (Fig. S6). However, despite the excellent separation, we note that HPLC requires specialized equipment. Moreover, HPLC purification results in high sample dilution, making this method less appropriate for small sample volumes, which is typically the case when primary, more expensive, antibodies are labeled. To address this limitation, we evaluated a second, size-based, purification technique using ultrafiltration. We performed a repetitive dilution concentration process using a regenerated cellulose membrane with a 100 kDa molecular weight cutoff (MWCO). In this process the large pG-ODN-antibody should be retained by the membrane, whereas the smaller pG-ODN should flow through. In contrast to SEC, ultrafiltration resulted in limited separation of the pG-ODN-antibody and pG-ODN with a recovery yield of 25% and contamination of 19% (Fig. 2d, bottom). The recovery yield and contamination percentage were calculated by gel analysis of purified product and known concentrations of a reference antibody (Fig. S7). Since pG-ODN is able to flow through the filter when applied in the absence of an antibody, we hypothesized that the limited separation was a result of the native affinity of pG for the antibody (Fig. S8). Taken together, these results show that the use of different (denaturing) washing buffers did improve the recovery yield (68%), however the separation remained limited.

In order to design a universal bench-top compatible purification method we therefore decided to turn our attention to antibody-specific purification methods. In addition to protein G, several recombinant proteins are available that target specific regions of an antibody, including protein A and L. While protein A overlaps with the binding site of protein G, protein L specifically binds to kappa light chains of mouse and human IgG monoclonal antibodies.^30^ Using commercially available magnetic beads functionalized with protein L, we were able to successfully purify pG-ODN functionalized Cetuximab (hIgG1) from pG-ODN (Fig. S9). However, the limited compatibility of protein L with different subclasses or host species of antibodies prevents the general application of this purification method. Therefore, we developed a purification method that utilizes the protein L magnetic beads to target free pG-ODN, instead of the labeled antibody. We achieved this *via* the covalent attachment of a scavenging antibody to protein L beads by crosslinking the primary amines of both proteins using a bis-succinimide ester-activated PEG compound BS(PEG)_5_ (Fig. 2d, top). Protein L ensures correct orientation of the scavenger antibody, making the Fc region available for free pG-ODN binding. We systematically optimized the cross-linking conditions, ensuring maximum capacity of the scavenger beads, and showed successful capture of pG-ODN (Fig. S10). Subsequently, we confirmed that this method can be used to purify pG-ODN labeled antibodies with a recovery of 58% and contamination of only 3% (Fig. 2d, bottom, S11). However, we note that the recovery yield is decreased to 30% when antibodies that have a high intrinsic affinity for protein L (hIgG, mIgG) are purified (Fig. S12). Additionally, it was shown that this scavenging approach could be performed on a small scale (5 µg) and was compatible with Fc-fusion proteins (Fig. S13). Protein L-functionalized magnetic beads, in combination with a scavenging antibody, therefore provide a universal platform for the small-scale purification of antibodies from different host species and sub-types.

To test the activity of the pG-ODN functionalized antibodies after labeling and purification we used fluorescent-activated cell sorting (FACS) analysis. First, we pre-incubated pG-ODN labeled Cetuximab that was purified either using SEC, ultrafiltration or magnetic scavenging beads with a CY5-functionalized complementary imager. Subsequently, we incubated A431 carcinoma cells, expressing high levels of EGFR, with pG-ODN-Cetuximab, hybridized to a CY5-labeled imager strand and subjected them to flow cytometry. In all cases an increase in the fluorescent intensity was observed, indicating binding of pG-ODN-Cetuximab to the EGFR receptor (Fig. 3a, S14).

**Figure 3.**
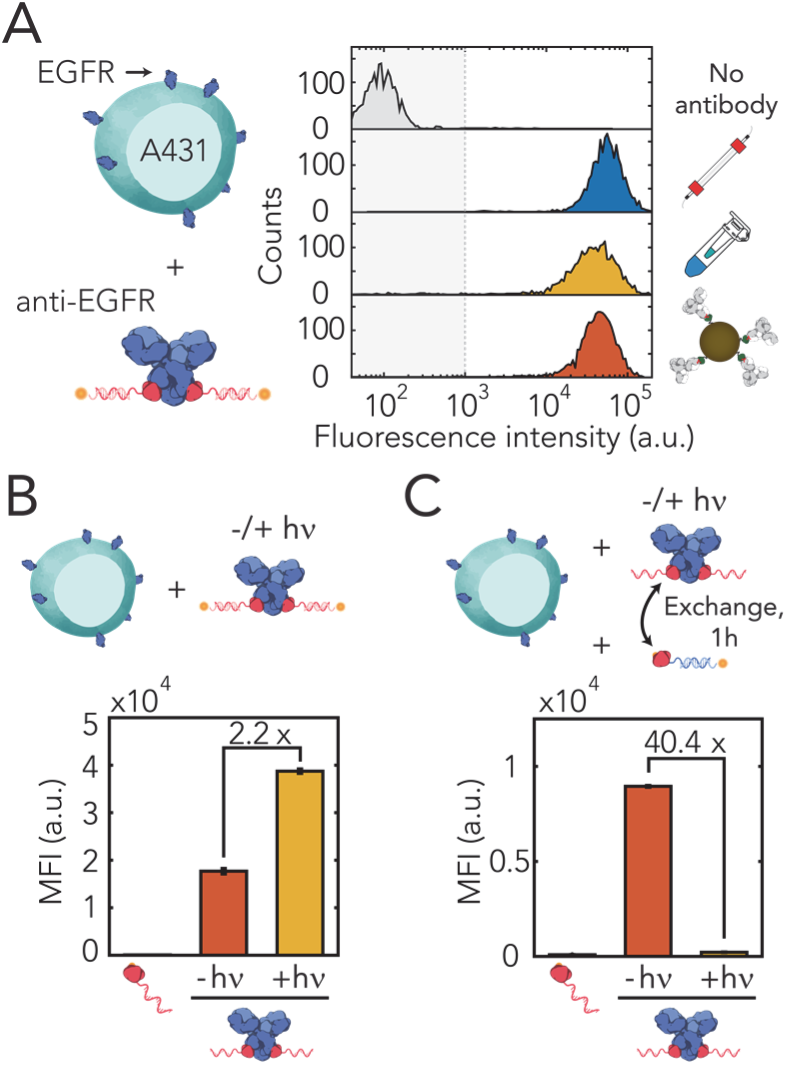
Flow cytometric analysis of EGFR-expressing A431 cells using 10 nM pG-ODN-functionalized Cetuximab hybridized to a CY5-labeled imager strand. (A) Antibody activity after purification using SEC, ultrafiltration and magnetic beads. Fluorescent intensity of pG-ODN-Cetuximab labeled A431 cells was compared to A431 cells incubated with only pG-ODN. (B) Labeling efficiency and (C) cross-contamination of pG-ODN non-covalently (-hν) or covalently (+hν) coupled to Cetuximab. pG-ODN-functionalized Cetuximab was incubated for 1h with 20 molar equivalents of a competing pG-ODN sequence. MFI represent the median fluorescence intensity and error bars represent SD (n=3).

To emphasize the importance of irreversible coupling, we compared the labeling efficiency and cross-contamination of antibodies labeled covalently (+hν) and non-covalently (-hν) with pG-ODN. To this end, Cetuximab was coupled to pG-ODN in the presence and absence of UV-illumination and directly incubated with A431 cells. After labeling, a complementary CY5-labeled imager strand was introduced and the cells were analyzed using flow cytometry. The 2.2-fold difference in median fluorescence shows a clear increase in labeling efficiency when the antibody is labeled covalently (Fig. 3b), which underlines the importance of irreversible labeling.

As cellular labeling procedures are typically performed with multiple antibodies, traces of pG-ODN present after purification could bind other antibodies resulting in unwanted cross-contamination. To evaluate the degree of cross-contamination we incubated non-covalently and covalently labeled pG-ODN-Cetuximab with a 20-fold molar excess of a competing pG-ODN docking sequence for 1h. Subsequently, A431 cells were incubated with the reaction mixture and after labeling, a CY5-functionalized imager strand, complementary to the competing pG-ODN sequence, was added. The observed median fluorescence for the cells that were labeled with non-covalent pG-ODN-Cetuximab showed pG-ODN exchange resulting in cross-contamination (Fig. 3c). In contrast, when pG-ODN was coupled covalently to Cetuximab *via* photo-induced coupling, no cross-contamination was observed. Additionally, it was shown that free Fc binding sites could be blocked during cellular labeling using 5-fold molar excess of pG compared to the competing pG-ODN sequence, nullifying cross-contamination (Fig. S15). These results show that multiple pG-ODN-functionalized antibodies can be used simultaneously, even when the pG-ODN antibody coupling is not quantitative and free Fc sites are still available.

Thus far, we have shown the synthesis and purification of pG-ODN-antibody constructs and used them successfully for cellular labeling. Next, we tested the multiplex applicability of the pG-ODN labeling strategy by coupling three primary antibodies, Cetuximab, mouse IgG2a anti-CD45 and anti-CD31, to a unique pG-ODN docking sequence. We performed simultaneous labeling of three cell types including A431 carcinoma cells, Jurkat T cells and Human Umbilical Vein Endothelial Cells (HUVECs) expressing EGFR, CD45 and CD31, respectively. Since quantitative pG-ODN labeling was not observed for all antibodies, we introduced a 5-fold molar excess of pG with respect to pG-ODN to block any remaining free Fc sites. After cellular labeling, we introduced a CY5-functionalized imager strand, complementary to a specific pG-ODN docking sequence, and analyzed the cells using flow cytometry (Fig. 4a). The presence of only one distinct fluorescent population per imager strand showed successful labeling of targets without exchange of pG-ODN sequences, verifying that pG-ODN-antibody conjugates are suitable for multiplexed cellular labeling and imaging (Fig. 4b).

**Figure 4.**
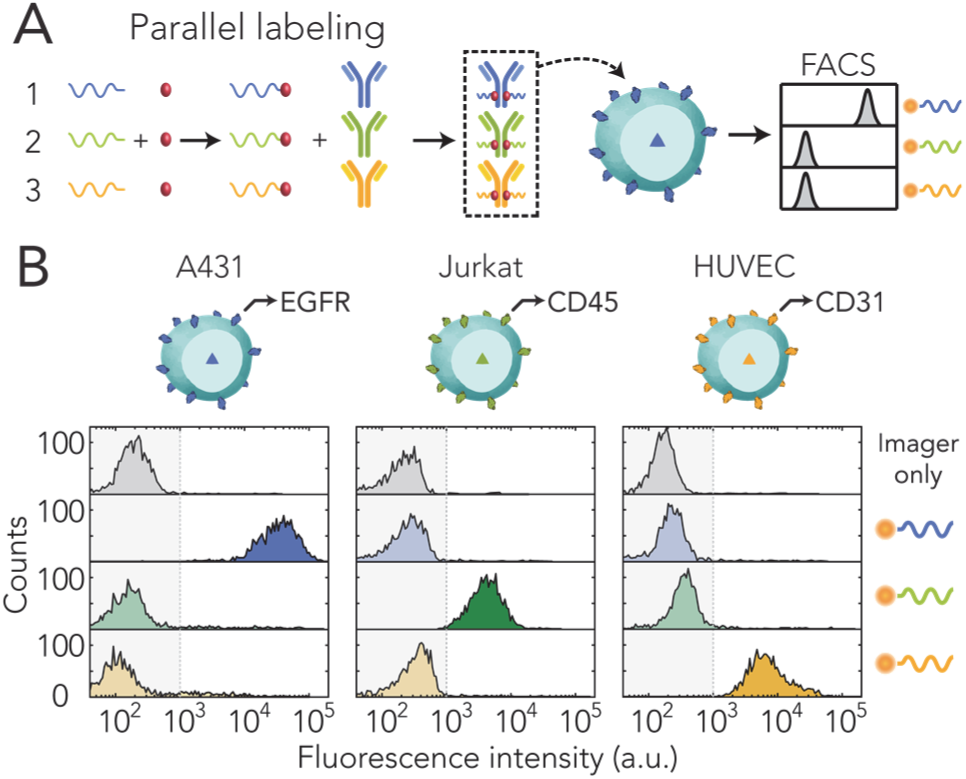
Characterization of protein G-ODN cross-talk. (A) Multiple antibodies are labeled in parallel with a unique pG-ODN sequence and incubated with cells expressing a specific target protein. Subsequently, CY5-labeled imager strands, complementary to a unique pG-ODN sequence, are used to detect the presence of each antibody on the cell surface. (B) Flow cytometric analysis of 3 cell lines (A431, Jurkat T cells and HUVECs) expressing EGFR, CD45 and CD31, respectively. Cells were incubated with 10 nM pG-ODN-labeled Cetuximab, mouse IgG2a anti-CD45 and mouse IgG2a anti-CD31, and subsequently labeled with 100 nM of a CY5-functionalized imager strand.

Although flow cytometry is an excellent tool to study cell populations, it is unable to provide a detailed view on the structure and organization of subcellular components. Super-resolution techniques, however, are able to visualize cellular targets with nanometer precision and therefore has the potential to increase our understanding of cell structure and function. DNA-PAINT and, in particular, exchange-PAINT rely on the use of antibodies functionalized with a unique ODN to achieve multiplexed super-resolution imaging with a spatial resolution down to ∼5 nm.^2,31^ To illustrate that pG-ODN-functionalized primary antibodies could directly be used for DNA-PAINT, we demonstrated *in situ* imaging in a fixed A431 carcinoma cell using pG-ODN-functionalized Cetuximab (Fig. 5a). We obtained super-resolution images of the EGFR receptor and observed little nonspecific binding when a non-complementary imager strand was used (Fig. 5b, S16). Recently, Jungmann and co-workers reported similar results when using ODN-functionalized protein A and G variants as secondary, non-covalent, labeling reagents in DNA-PAINT.^32^ Since pG-ODN is covalently coupled to the antibody using UV light, we additionally investigated if the pG-ODN antibody coupling is reversed when the construct is exposed to high laser intensity. To validate the stability of the pGODN-antibody construct we quantified the number of localizations during image acquisition over the course of 8 minutes (Fig. 5c). The observed number of localizations remained constant over time, confirming that the photo-conjugated pG-ODN-antibody construct is stable when exposed to high laser intensity. This result also allowed the use of pG-ODN-antibody constructs for cellular staining in combination with direct STORM (dSTORM), which requires high illumination power. Successful dSTORM images of the EGFR receptor on A431 cells were acquired using a complementary CY5-functionalized imager strand, hybridized to a pG-ODN-Cetuximab construct (Fig. 5d, S17). Taken together, these results show the compatibility of the pG-ODN labeling approach with super-resolution techniques, including DNA-PAINT and dSTORM.

**Figure 5.**
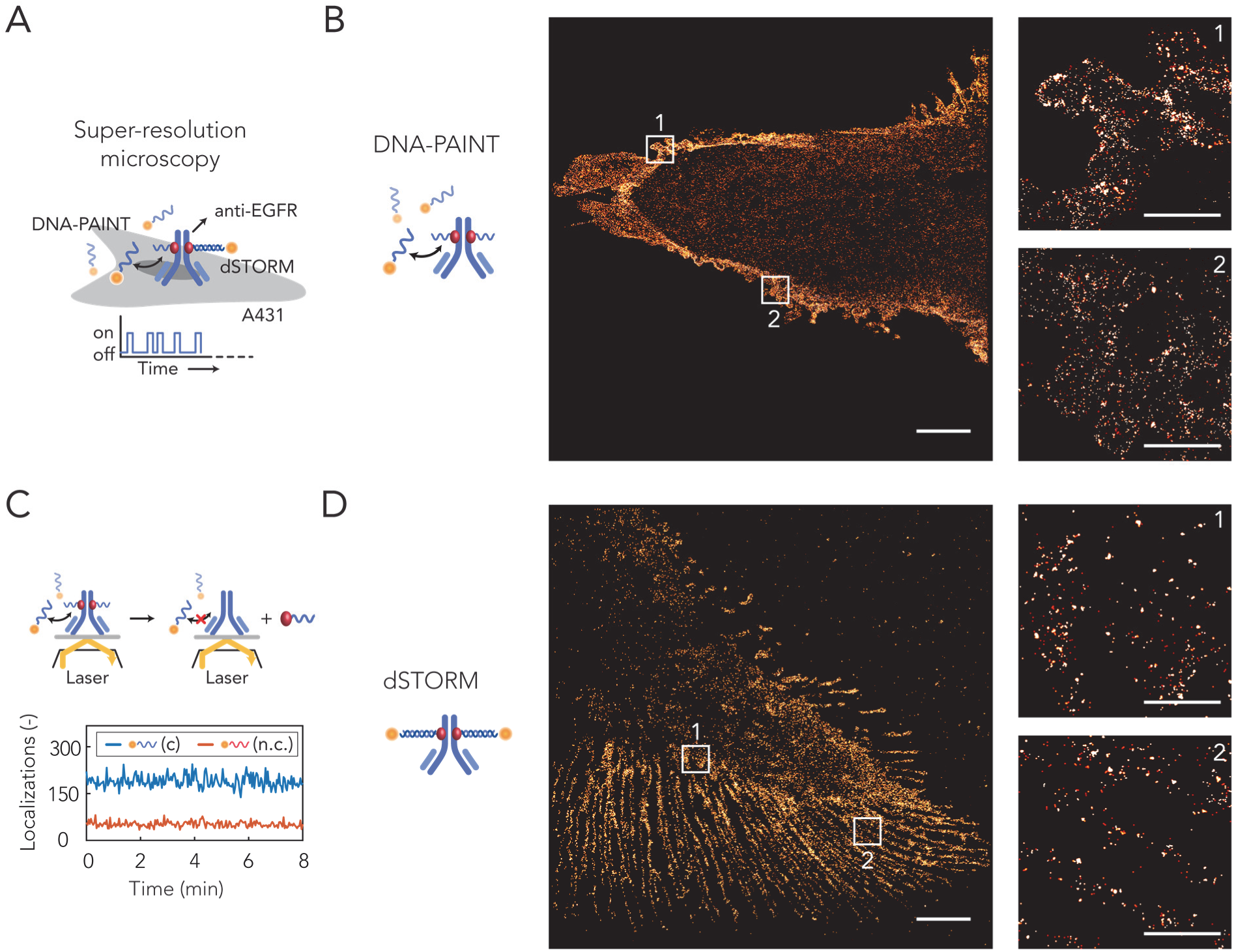
Protein G-ODN-antibody conjugates for applications in super-resolution microscopy. (A) A431 carcinoma cells are labeled with a pG-ODN-Cetuximab construct containing either a short (11 nt) or a long (20 nt) docking strand. Subsequently, the labeled cells are fixated to a glass slide and super-resolution images are obtained using the transient binding of a short imager strand (DNA-PAINT) or using a long irreversible binder in combination with a photoswitchable dye (dSTORM). (B) DNA-PAINT super-resolution image obtained using ATTO647N-functionalized imager strands (20,000 frames, 20-Hz frame rate). Zoom in on 2 defined areas show the distribution of EGFR receptors (C) Number of localizations observed over the course of 8 minutes during a DNA-PAINT acquisition using complementary (c) or non-complementary (n.c.) imager strands. (D) Super-resolution image obtained with dSTORM using a CY5-functionalized imager strand (20,000 frames, 65.5-Hz frame rate), including a zoom in on 2 defined areas which show the distribution of EGFR receptors. Scale bars, 5 µm and 1 µm for the zoom in images.

## Conclusion

In this work, we have developed a generally applicable ODN-antibody coupling method using a small protein G adaptor that site-specifically targets the Fc domain of an IgG antibody. We successfully demonstrated pG-ODN labeling of antibodies from different host species without cross-reactivity towards other proteins. Importantly, we showed that the pG-ODN labeling did not affect the native function of the antibody. In combination with the universal, benchtop-compatible, purification strategy using magnetic beads this ODN labeling method is directly applicable to commercially available primary antibodies. Since multiple pG-ODN conjugates can be constructed in parallel and lyophilized without loss of function, the potential of the pG-ODN labeling strategy lies in the synthesis of a library of pG-ODN constructs that can directly be used for antibody labeling. This could eventually facilitate the implementation of the multiplexing abilities of DNA based read-out methods for the detection of a large variety of subcellular components and make these methods accessible for a broader scientific community. Additionally, we envision the use of pG-ODN-antibody constructs in more quantitative imaging applications, owing to the unique ability of pG-ODN conjugates to specifically label an antibody with a controlled number of ODNs.

## Materials and Methods

### Recombinant protein cloning, expression and purification

A pET28a(+) vector encoding the pG gene as reported previously26 was site-specifically mutated by the insertion of an N-terminal serine using the QuikChange Lightning Multi Site-Directed Mutagenesis kit (Agilent), according to the manufacturer’s instructions using following forward and reverse primers (ser codon underlined): 5’-TTTAAGAAGGAGATATAACATG<u>AGT</u>TGCTGGTCCCATCCG-3’ and 5’-CGGATGGGACCAGCA<u>ACT</u>CATGTTATATC TCCTTCTTAAAG-3’, respectively. Insertion was confirmed by DNA sequencing service provided by StarSEQ® (Mainz, Germany). Plasmid DNA of the pG gene was co-transformed with the pEVOL-pBpF vector, encoding for the orthogonal aminoacyl tRNA synthetase/tRNA pair (kindly provided by Peter Schultz), into *E. coli* BL21(DE3) (Novagen) for protein expression. A single colony of freshly transformed cells was cultured at 37 °C in 500 mL 2xYT medium supplemented with 50 µg/mL kanamycin (Merck) and 25 µg/mL chloramphenicol (Sigma Aldrich). When the OD_600_ of the culture reached ∼0.6 protein expression was induced by addition of β-D-1-thiogalactopyranoside (IPTG, Applichem), arabinose (Sigma Aldrich) and the unnatural amino acid *p-*BpA (Bachem) in a final concentration of 1 mM, 0.02% (w/v) and 1 mM, respectively. The induced protein expression was carried out for ∼18h at 25 °C and subsequently the cells were harvested by centrifugation at 10,000 xg for 10 min at 4 °C. The cell pellet was resuspended in Bug-Buster (5 mL/g pellet, Merck) supplemented with benzonase 5 µL/g pellet, Merck) and incubated for 45 minutes on a shaking table. The suspension was centrifugated at 40,000 xg for 30 min at 4 °C and the supernatant was subjected to Ni-NTA affinity chromatography on a gravity column. His-tagged pG was loaded on the column and washed with washing buffer (1x PBS, 370 mM NaCl, 10% (v/v) glycerol, 20 mM imidazole, pH 7.4) before elution with his-elution buffer (1x PBS, 370 mM NaCl, 10% (v/v) glycerol, 250 mM imidazole, pH 7.4). Subsequently, the Ni-NTA elution fraction was loaded on a *Strep*-tactin column, washed with washing buffer (100 mM Tris-HCl, 150 mM NaCl, 1 mM EDTA, pH 8.0) and eluted using wash buffer supplemented with 2.5 mM desthiobiotin (IBA Life Sciences). Proteins were stored at −80 °C in 1 mL aliquots of 50 µM at −80°C in a buffer containing 100 mM Tris-HCl, 150 mM NaCl, 1 mM EDTA and 2 mM TCEP at pH 8.0. The concentration of pG was calculated on the basis of the absorption at 280 nm (ND-1000, Thermo Scientific) assuming an extinction coefficient of 15,470 M-1 cm-1 and the purity of pG was assessed on reducing SDS-PAGE and liquid chromatography quadrupole time-of-flight mass spectrometry (Q-Tof).

### Preparation of reaction ODNs

In a typical reaction, to a solution of 10 nmol ODN in water (10 µL) was added 1x PBS, pH 7.2 (30 µL) and 100 nmol Sulfo-SMCC (Thermo scientific) in DMSO (40 µL). The reaction was incubated at 850 RPM for 2h at 20 °C. Excess Sulfo-SMCC was removed using 2 rounds of ethanol precipitation. SMCC-labeled ODNS were precipitated by the addition of 10% (v/v) 5 M NaCl and 300% (v/v) ice-cold EtOH and incubating for 75 minutes at −30 °C. The reaction mixture was centrifuged at 19,000 xg for 30 min at 4°C, the pellet was reconstituted in 1x PBS (pH 7.2) and the precipitation was repeated. After centrifugation the pellet was washed in 95% (v/v, in water) ice-cold EtOH, centrifuged at 19,000 xg for 15 min and lyophilized.

### General procedure for conjugation of ODN to pG

For conjugation of pG to a SMCC-functionalized ODN an aliquot of pG was buffer exchanged to (100 mM Sodium Phosphate, 25 µM TCEP, pH 7) using a PD10 desalting column (GE Healthcare). Subsequently, desalted pG was concentrated to a final concentration of 50 µM using Amicon 3 kDa MWCO centrifugal filters (Merck Millipore). 10 nmol lyophilized SMCC-functionalized ODN was reconstituted in 40 µL 50 µM pG (2 nmol) resulting in a 5x excess of maleimide-ODN. The reaction was shaken at 850 RPM for 3h at 20 °C. The coupling efficiency was assessed using SDS-PAGE under non-reducing conditions. Purification, if applicable, of pG-ODN was performed using fast protein liquid chromatography (FPLC, ÄKTA Prime, GE Healthcare) with an anion-exchange HiTrap Q HP column (1 mL, GE Healthcare) using a salt gradient with a start and end concentration of 100 and 500 mM NaCl in 50 mM Tris-HCl (pH 7.5), respectively. Elution fractions were collected and analyzed by measuring on-line absorption at 280 nm and SDS-PAGE under non-reducing conditions. pG-ODN conjugates were aliquoted and stored at −80 °C.

### General procedure for pG-ODN antibody labeling

Before conjugation of the antibody to the pG-ODN, all antibodies are buffer exchanged to 1x PBS (pH 7.4) using Amicon 10 kDa MWCO centrifugal filters (Merck Millipore).

#### In the absence of BSA

In a typical reaction for SDS-PAGE gel analysis, a 10 µL aliquot containing 0.4 µM of antibody and 4 µM of pG-ODN is exposed for 1h to UV light (λ = 365 nm) at 4 °C.

#### In the presence of BSA

For the conjugation of the antibody to pG-ODN in the presence of BSA the pG-ODN reaction mixture was first incubated with 15 nmol of DTT and shaken at 850 RPM for 30 minutes at 20 °C to deactivate remaining maleimide-ODNs. Subsequently, pG-ODN was buffer exchanged using a Zeba(tm) spin desalting column, 7000 MWCO, 0.5 mL (Thermo scientific).

### General procedure for antibody functionalization of protein L magnetic beads

The scavenging antibody Cetuximab (Erbitux, Merck) was buffer exchanged to conjugation buffer (100 mM Sodium Phosphate, 70 mM NaCl, 0.05% TWEEN-20 (v/v), pH 8.0) using a Zeba(tm) spin desalting column, 7000 MWCO, 0.5 mL (Thermo scientific) according to the manufacturer’s instructions. In a typical reaction 25 µL of Pierce(tm) protein L magnetic beads was added to 75 µL conjugation buffer. The beads were washed twice in 200 µL conjugation buffer after which the magnetic beads were incubated with 50 µg scavenging antibody in a final volume of 250 µL. The tube was rotated for 1h at room temperature. To crosslink the antibody to the beads 0.5 µL of 250 mM BS(PEG)5 (Thermo scientific) (dissolved in dry DMSO) was added to the reaction mixture. Crosslinking was performed for 30 minutes in a rotating wheel at room temperature, after which the reaction was deactivated by the addition of 25 µL 1 M Tris-HCl (pH 7.5). Deactivation was performed for 15 min in a rotating wheel at room temperature. Subsequently, the magnetic beads were collected with a magnetic stand and the supernatant was discarded. The scavenging beads were incubated 2 times for 5 minutes with 200 µL washing buffer (100 mM Tris-HCl, 1 M NaSCN, pH 7.5), subsequently washed 2 times with 200 µL storage buffer (1x PBS, 0.05% TWEEN-20 (v/v), pH 7.4) and stored at 4 °C.

### General procedure for purification of pG-ODN functionalized antibody

This procedure is optimized to purify µL of 4 µM antibody-pG-ODN from 40 µM uncoupled pG-ODN using 25 µL of magnetic scavenging beads. 25 µL of scavenging beads were washed two times in binding buffer (1x PBS, 870 mM NaCl, 0.05% TWEEN-20 (v/v), pH 7.4). The beads were redissolved in 37.5 µL binding buffer and 12.5 µL of the antibody/pG-ODN reaction mixture was added and incubated for 15 minutes in a rotating wheel at room temperature. The scavenging beads were collected with a magnetic stand and the supernatant was discarded. Subsequently, the beads were incubated 2 times for 5 minutes with 200 µL washing buffer (100 mM Tris-HCl, 1 M NaSCN, pH 7.5). The whole process was repeated two more times to remove all pG-ODN. Amicon 50 kDa MWCO centrifugal filters were used to purify the anti-body-pG-ODN conjugates from unreacted maleimide-ODN conjugates and buffer-exchange the purified conjugates to 1x PBS, pH 7.4. Removal of the pG-ODN conjugate and recovery of the antibody coupled pG-ODN were assessed based on gel band intensity analysis using SDS-PAGE under reducing conditions.

### Cellular labeling for flow cytometry

A431, Jurkat T and HUVEC cells were cultured in a 175 cm2 flask. Cells were harvested at a confluency of ∼80%, washed in labeling buffer (1x PBS, 0.1% BSA (w/v), pH 7.4) and diluted to a final concentration of 3.5*106 cells/mL in labeling buffer. Subsequently, 12.5 µL of the cell suspension was incubated in a final volume of 250 µL labeling buffer containing 10 nM of pG-ODN labeled antibody. The reaction mixture was shaken at 400 RPM for 30 minutes at room temperature. Subsequently, the labeled cells were centrifuged for 5 minutes at 150 xg and the supernatant was removed. The pelleted cells were redissolved in labeling buffer containing 100 nM of the complementary CY5-labeled ODN. The reaction mixture was incubated and centrifuged as described above and subsequently analyzed using flow cytometry. For the multiplexed experiment (Figure 4) the labeling buffer was supplemented with 1.5 µM pG.

### Super Resolution Sample Preparation and Imaging

A431 cells (ATCC®CRL-1555(™)) were seeded on an 8 well glass bottom μ-slide (ibidi GmbH, Germany) overnight. Live-cell immunolabelling started with 5 minutes acclimation of the cells at room temperature, followed by 5 minutes acclimation at 4°C. Cells were incubated for 45 minutes at 4°C in DMEM 3% BSA with 1 μg/mL of pG-ODN-Cetuximab. Then, three 5 minutes washes with PBS at 4°C were followed by formalin 5% and glutaraldehyde 0.25% fixation for 10 minutes. Ultimately, cells were washed three times for 5 minutes with PBS and stored in the fridge before imaging. Super resolution images were acquired using a Nikon N-STORM in total internal reflection fluorescence (TIRF) mode. The system is equipped with a Nikon 100x, 1.49 NA oil immersion objective and an Andor iXON3 camera. Images were acquired onto a 256 x 256 pixel region (40.96 x 40.96 μm) and analyzed with NIS Element Nikon software. Samples were illuminated with a 647 nm laser at 140 mW, 16 ms and 20.000 frames for STORM or 70 mW, 50 ms and 20.000 frames for DNA-PAINT. To perform STORM imaging we used a specific buffer to induce dye photoswitching: 5% w/v glucose, 100 mM cysteamine, 0.5 mg/mL glucose oxidase and 40 µg/mL catalase in PBS. For DNA-PAINT imaging we diluted DNA imager-Atto647N to 1nM in PBS 500 mM NaCl.

## ASSOCIATED CONTENT

## ACKNOWLEDGMENTS

We thank L.A. Tiemeijer & C.M. Sahlgren for providing and culturing of the HUVEC cells. Sinem K. Saka from the Wyss Institute for Biologically Inspired Engineering is gratefully acknowledged for valuable insights and fruitful discussions. This work was supported by the European Research Council, ERC (project n. 677313 BioCircuit) an NWO-VIDI grant from the Netherlands Organization for Scientific Research (NWO, 723.016.003), funding from the Ministry of Education, Culture and Science (Gravity programs, 024.001.035 & 024.003.013)

---

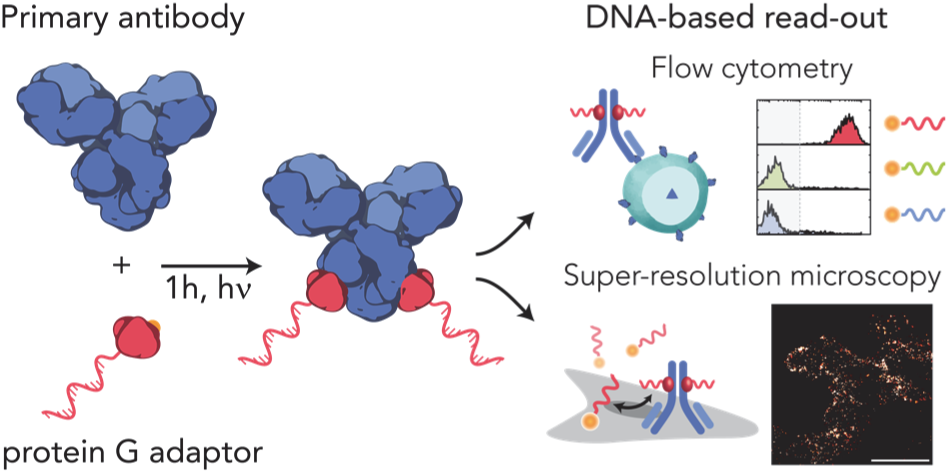

## REFERENCES

(1) Chen, Y.-J.; Groves, B.; Muscat, R. A.; Seelig, G. DNA Nanotechnology from the Test Tube to the Cell. Nat. Nanotechnol. 2015, 10 (9), 748–760. https://doi.org/10.1038/nnano.2015.195.

(2) Jungmann, R.; Avendaño, M. S.; Woehrstein, J. B.; Dai, M.; Shih, W. M.; Yin, P. Multiplexed 3D Cellular Super-Resolution Imaging with DNA-PAINT and Exchange-PAINT. Nat. Methods 2014, 11 (3), 313–318. https://doi.org/10.1038/nmeth.2835.

(3) Wade, O. K.; Woehrstein, J. B.; Nickels, P. C.; Strauss, S.; Stehr, F.; Stein, J.; Schueder, F.; Strauss, M. T.; Ganji, M.; Schnitzbauer, J.; et al. 124-Color Super-Resolution Imaging by Engineering DNA-PAINT Blinking Kinetics. Nano Lett. 2019. https://doi.org/10.1021/acs.nanolett.9b00508.

(4) Jungmann, R.; Steinhauer, C.; Scheible, M.; Kuzyk, A.; Tinnefeld, P.; Simmel, F. C. Single-Molecule Kinetics and Super-Resolution Microscopy by Fluorescence Imaging of Transient Binding on DNA Origami. Nano Lett. 2010, 10 (11), 4756–4761. https://doi.org/10.1021/nl103427w.

(5) Chen, K. H.; Boettiger, A. N.; Moffitt, J. R.; Wang, S.; Zhuang, X. Spatially Resolved, Highly Multiplexed RNA Profiling in Single Cells. Science 2015, 348 (6233), aaa6090–aaa6090. https://doi.org/10.1126/science.aaa6090.

(6) Fredriksson, S.; Gullberg, M.; Jarvius, J.; Olsson, C.; Pietras, K.; Gústafsdóttir, S. M.; Östman, A.; Landegren, U. Protein Detection Using Proximity-Dependent DNA Ligation Assays. Nat. Biotechnol. 2002, 20 (5), 473–477. https://doi.org/10.1038/nbt0502-473.

(7) Sano, T.; Smith, C. L.; Cantor, C. R. Immuno-PCR: Very Sensitive Antigen Detection by Means of Specific Antibody-DNA Conjugates. Science 1992, 258 (5079,), 120–122.

(8) Bailey, R. C.; Kwong, G. A.; Radu, C. G.; Witte, O. N.; Heath, J. R. DNA-Encoded Antibody Libraries: A Unified Platform for Multiplexed Cell Sorting and Detection of Genes and Proteins. J. Am. Chem. Soc. 2007, 129 (7), 1959–1967. https://doi.org/10.1021/ja065930i.

(9) Dahotre, S. N.; Chang, Y. M.; Wieland, A.; Stammen, S. R.; Kwong, G. A. Individually Addressable and Dynamic DNA Gates for Multiplexed Cell Sorting. Proc. Natl. Acad. Sci. 2018, 115 (17), 4357–4362. https://doi.org/10.1073/pnas.1714820115.

(10) Tam, J.; Cordier, G. A.; Borbely, J. S.; Sandoval Álvarez, Á.; Lakadamyali, M. Cross-Talk-Free Multi-Color STORM Imaging Using a Single Fluorophore. PLoS ONE 2014, 9 (7), e101772. https://doi.org/10.1371/journal.pone.0101772.

(11) Rothemund, P. W. K. Folding DNA to Create Nanoscale Shapes and Patterns. Nature 2006, 440 (7082), 297–302. https://doi.org/10.1038/nature04586.

(12) Dietz, H.; Douglas, S. M.; Shih, W. M. Folding DNA into Twisted and Curved Nanoscale Shapes. Science 2009, 325 (5941), 725–730. https://doi.org/10.1126/science.1174251.

(13) Douglas, S. M.; Dietz, H.; Liedl, T.; Högberg, B.; Graf, F.; Shih, W. M. Self-Assembly of DNA into Nanoscale Three-Dimensional Shapes. Nature 2009, 459 (7245), 414–418. https://doi.org/10.1038/nature08016.

(14) Lin, C.; Jungmann, R.; Leifer, A. M.; Li, C.; Levner, D.; Church, G. M.; Shih, W. M.; Yin, P. Submicrometre Geometrically Encoded Fluorescent Barcodes Self-Assembled from DNA. Nat. Chem. 2012, 4 (10), 832–839. https://doi.org/10.1038/nchem.1451.

(15) Woehrstein, J. B.; Strauss, M. T.; Ong, L. L.; Wei, B.; Zhang, D. Y.; Jungmann, R.; Yin, P. Sub–100-Nm Metafluorophores with Digitally Tunable Optical Properties Self-Assembled from DNA. Sci. Adv. 2017, 3 (6), e1602128. https://doi.org/10.1126/sciadv.1602128.

(16) Agasti, S. S.; Wang, Y.; Schueder, F.; Sukumar, A.; Jungmann, R.; Yin, P. DNA-Barcoded Labeling Probes for Highly Multiplexed Exchange-PAINT Imaging. Chem. Sci. 2017, 8 (4), 3080–3091. https://doi.org/10.1039/C6SC05420J.

(17) Trads, J. B.; Tørring, T.; Gothelf, K. V. Site-Selective Conjugation of Native Proteins with DNA. Acc. Chem. Res. 2017, 50 (6), 1367–1374. https://doi.org/10.1021/acs.accounts.6b00618.

(18) Szabó, Á. The Effect of Fluorophore Conjugation on Antibody Affinity and the Photophysical Properties of Dyes. Biophys. J. 2018, 114 (3), 688–700.

(19) Kazane, S. A.; Sok, D.; Cho, E. H.; Uson, M. L.; Kuhn, P.; Schultz, P. G.; Smider, V. V. Site-Specific DNA-Antibody Conjugates for Specific and Sensitive Immuno-PCR. Proc. Natl. Acad. Sci. 2012, 109 (10), 3731–3736. https://doi.org/10.1073/pnas.1120682109.

(20) Keppler, A.; Gendreizig, S.; Gronemeyer, T.; Pick, H.; Vogel, H.; Johnsson, K. A General Method for the Covalent Labeling of Fusion Proteins with Small Molecules in Vivo. Nat. Biotechnol. 2003, 21 (1), 86–89. https://doi.org/10.1038/nbt765.

(21) Los, G. V.; Encell, L. P.; McDougall, M. G.; Hartzell, D. D.; Karassina, N.; Zimprich, C.; Wood, M. G.; Learish, R.; Ohana, R. F.; Urh, M.; et al. HaloTag: A Novel Protein Labeling Technology for Cell Imaging and Protein Analysis. ACS Chem. Biol. 2008, 3 (6), 373–382. https://doi.org/10.1021/cb800025k.

(22) Gautier, A.; Juillerat, A.; Heinis, C.; Corrêa, I. R.; Kindermann, M.; Beaufils, F.; Johnsson, K. An Engineered Protein Tag for Multiprotein Labeling in Living Cells. Chem. Biol. 2008, 15 (2), 128–136. https://doi.org/10.1016/j.chembiol.2008.01.007.

(23) Rosen, C. B.; Kodal, A. L. B.; Nielsen, J. S.; Schaffert, D. H.; Scavenius, C.; Okholm, A. H.; Voigt, N. V.; Enghild, J. J.; Kjems, J.; Tørring, T.; et al. Template-Directed Covalent Conjugation of DNA to Native Antibodies, Transferrin and Other Metal-Binding Proteins. Nat. Chem. 2014, 6 (9), 804–809. https://doi.org/10.1038/nchem.2003.

(24) Yu, C.; Tang, J.; Loredo, A.; Chen, Y.; Jung, S. Y.; Jain, A.; Gordon, A.; Xiao, H. Proximity-Induced Site-Specific Antibody Conjugation. Bioconjug. Chem. 2018, 29 (11), 3522–3526. https://doi.org/10.1021/acs.bioconjchem.8b00680.

(25) Hui, J. Z.; Tamsen, S.; Song, Y.; Tsourkas, A. LASIC: Light Activated Site-Specific Conjugation of Native IgGs. Bioconjug. Chem. 2015, 26 (8), 1456–1460. https://doi.org/10.1021/acs.bio-conjchem.5b00275.

(26) Rosier, B. J. H. M.; Cremers, G. A. O.; Engelen, W.; Merkx, M.; Brunsveld, L.; de Greef, T. F. A. Incorporation of Native Antibodies and Fc-Fusion Proteins on DNA Nanostructures via a Modular Conjugation Strategy. Chem. Commun. 2017, 53 (53), 7393–7396. https://doi.org/10.1039/C7CC04178K.

(27) Antibodies, Proteins, Kits and Reagents for Life Science Abcam https://www.abcam.com/ (accessed Apr 23, 2019).

(28) Rust, M. J.; Bates, M.; Zhuang, X. Sub-Diffraction-Limit Imaging by Stochastic Optical Reconstruction Microscopy (STORM). Nat. Methods 2006, 3 (10), 793–796. https://doi.org/10.1038/nmeth929.

(29) Navaratnam, S.; Hamblett, I.; Tonnesen, H. H. Photoreactivity of Biologically Active Compounds. XVI. Formation and Reactivity of Free Radicals in Mefloquine. J. Photochem. Photobiol. B 2000, 56 (1), 25–38.

(30) Björck, L. Protein L. A Novel Bacterial Cell Wall Protein with Affinity for Ig L Chains. J. Immunol. Baltim. Md 1950 1988, 140 (4), 1194–1197.

(31) Schnitzbauer, J.; Strauss, M. T.; Schlichthaerle, T.; Schueder, F.; Jungmann, R. Super-Resolution Microscopy with DNA-PAINT. Nat. Protoc. 2017, 12 (6), 1198–1228. https://doi.org/10.1038/nprot.2017.024.

(32) Schlichthaerle, T.; Ganji, M.; Auer, A.; Wade, O. K.; Jungmann, R. Bacterial-Derived Antibody Binders as Small Adapters for DNA-PAINT Microscopy. ChemBioChem 2018. https://doi.org/10.1002/cbic.201800743.

