## Supporting Information for "Efficient small-scale conjugation of DNA to primary antibodies for multiplexed cellular targeting"

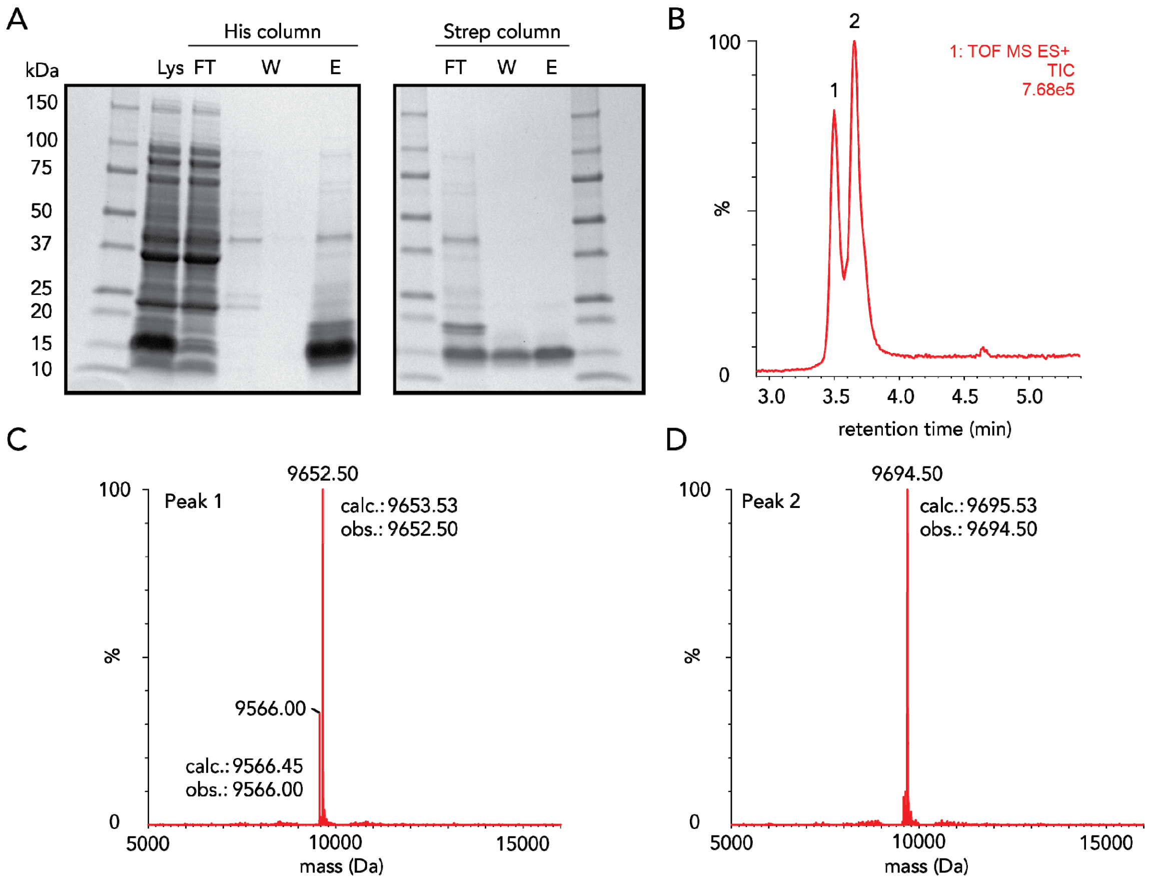


**Figure S1**. Purification and characterization of pG. (A) SDS-PAGE gel analysis of affinity-tag based purification of pG. (calculated mass: 9784.64 Da). pG runs at an apparent mass of 14 kDa. Labels: Lys, lysate; FT, flow through; W, wash; E, elution. (B) Chromatogram of pG analyzed by liquid chromatography quadrupole time-of-flight mass spectrometry. (C) Deconvoluted mass spectrum of peak 1 shows 2 peaks corresponding to pG without the N-terminal methionine (9652.5 Da) and serine (9566.0 Da). (D) Deconvoluted mass spectrum of peak 2 shows 1 peak corresponding to acetylated pG without the N-terminal methionine (9694.5 Da).

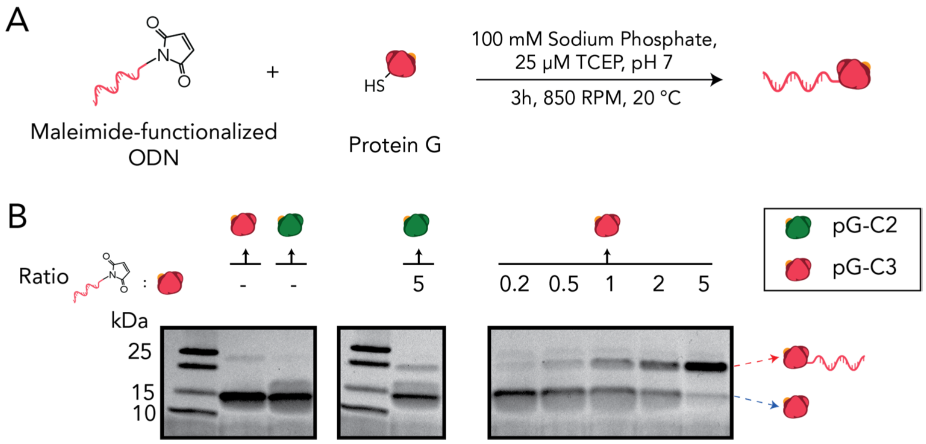


**Figure S2**. Schematic overview of the protein G-oligonucleotide (pG-ODN) coupling. (A) Reaction conditions for pG-ODN coupling. (B) SDS-PAGE gel analysis under non-reducing conditions of the pG-ODN reaction mixture. Using a 5-fold molar excess of maleimide-functionalized ODN, >90% of pG-C3 is successfully labeled with an ODN compared to ~15% of pG-C2. Labels: pG-C2, pG with the cysteine directly after the N-terminal methionine; pG-C3, pG with a serine placed N-terminally of the cysteine. Amino acid sequence of both pG constructs is shown in table 2.

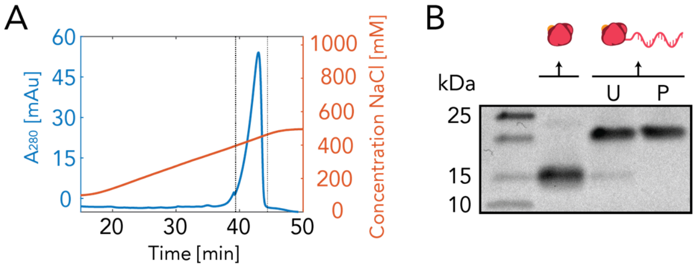


**Figure S3**. Overview of the purification of pG-ODN constructs. (A) Elution trace of pG-ODN using fast protein liquid chromatography (FPLC) monitored by on-line absorption at 280 nm. The large peak, indicated by the dashed lines, was collected. (B) SDS-PAGE gel analysis under non-reducing conditions of the collected fraction, showing successful removal of uncoupled pG. Labels: U, unpurified pG-ODN reaction mixture; P, purified pG-ODN

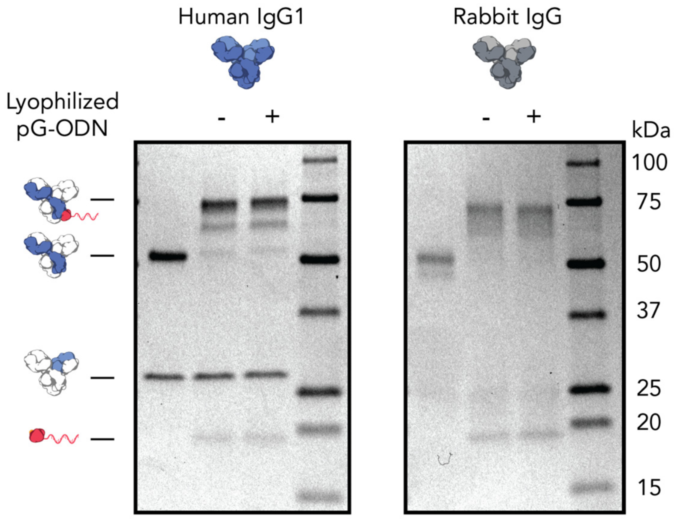


**Figure S4**. SDS-PAGE gel analysis under reducing conditions of pG-ODN antibody coupling before (-) and after (+) lyophilization of pG-ODN. For both Cetuximab (hIgG1) and Rabbit IgG, no decrease in coupling efficiency is observed when pG-ODN was lyophilized prior to antibody coupling.

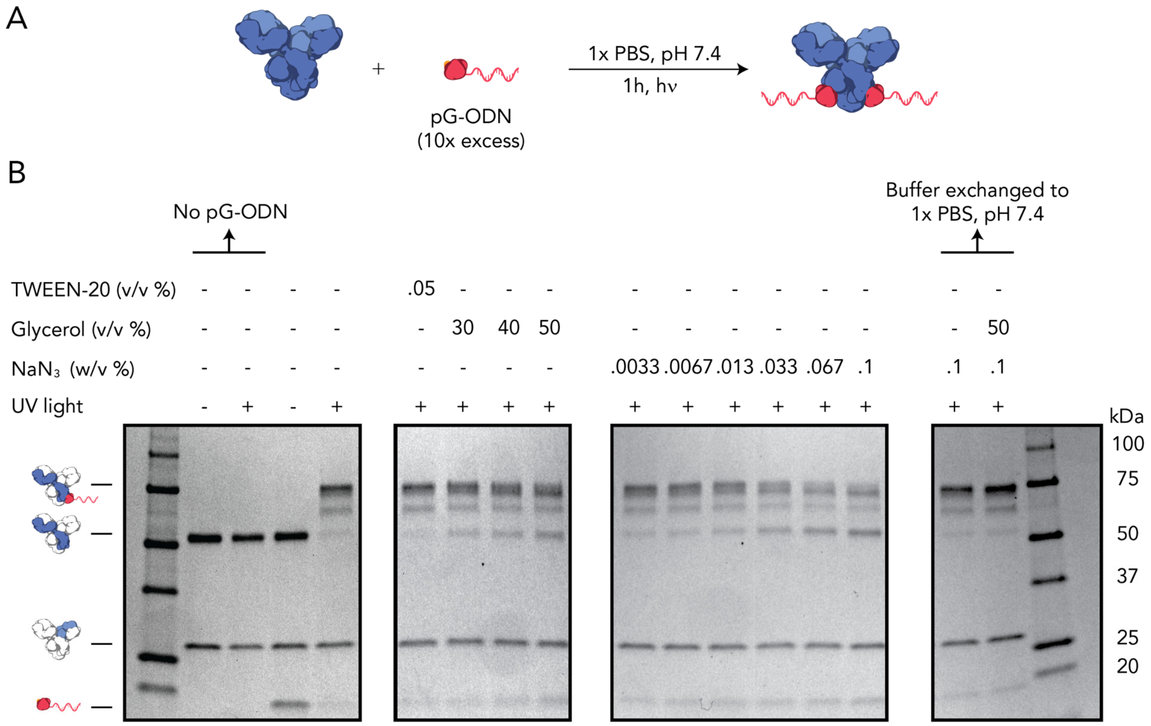


**Figure S5**. Coupling efficiency of pG-ODN to Cetuximab in the presence of multiple additives. (A) Schematic overview of the reaction conditions. (B) SDS-PAGE gel analysis under reducing conditions of Cetuximab coupled to pG in the presence of TWEEN-20, glycerol and sodium azide ($NaN_3$). Both glycerol and sodium azide inhibit pG-ODN-antibody formation. Buffer exchanging an antibody solution that contains 0.1% sodium azide in the presence and absence of 50% glycerol to 1x PBS, pH 7.4 using ultrafiltration fully restores the coupling efficiency.

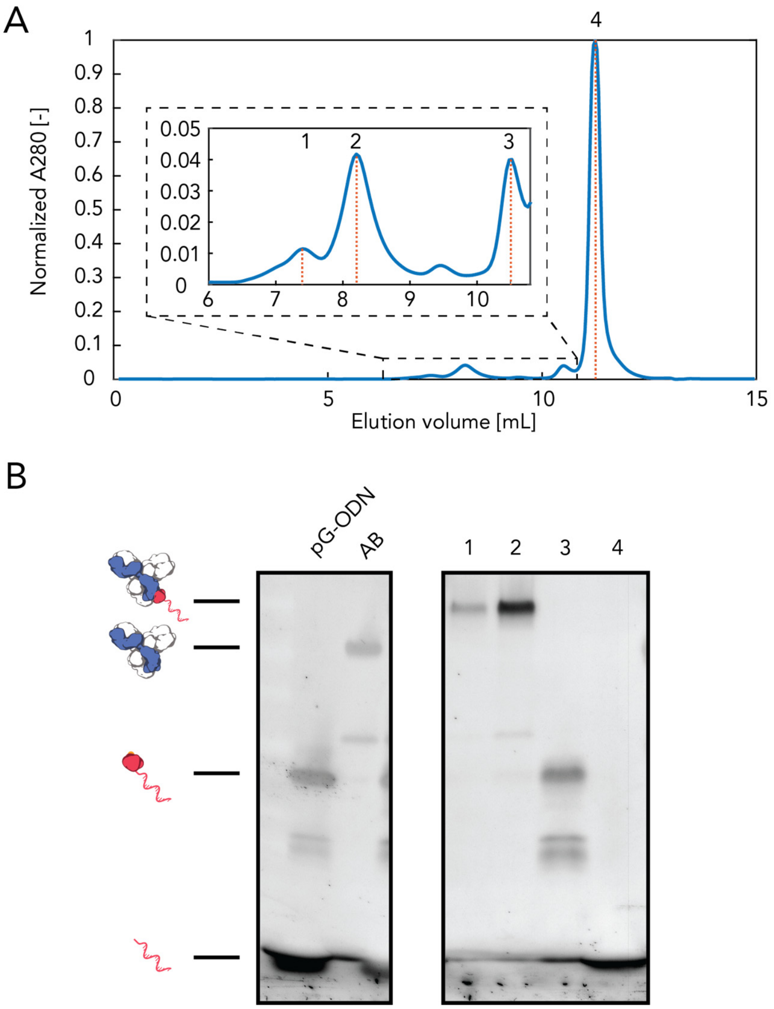


**Figure S6**. Purification of pG-ODN-antibody constructs using size exclusion chromatography (SEC). (A) Chromatogram showing the separation of pG-ODN-antibody (1, 2), pG-ODN (3) and the uncoupled ODN (4). (B) SDS-PAGE gel analysis of the 4 peaks shown in the chromatogram stained with SYBR Gold shows the successful separation of pG-ODN-antibody and pG-ODN constructs. Label: AB, Antibody (Cetuximab)

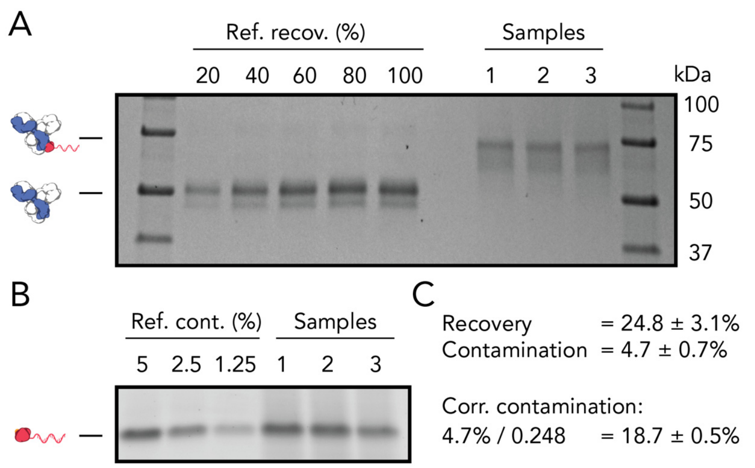


**Figure S7**. Recovery and purity of pG-ODN-antibody constructs purified by ultrafiltration. (A) SDS-PAGE gel analysis under reducing conditions stained with Coomassie Blue. The gel band intensity corresponding to the recovered heavy chain of the pG-ODN-antibody was compared to a reference. (B) SDS-PAGE gel analysis under non-reducing conditions stained with SYBR Gold. The intensity of the band corresponding to pG-ODN after purification was compared to a reference. (C) Calculation of the recovery and contamination. The contamination was corrected based on the recovery of the pG-ODN-antibody construct.

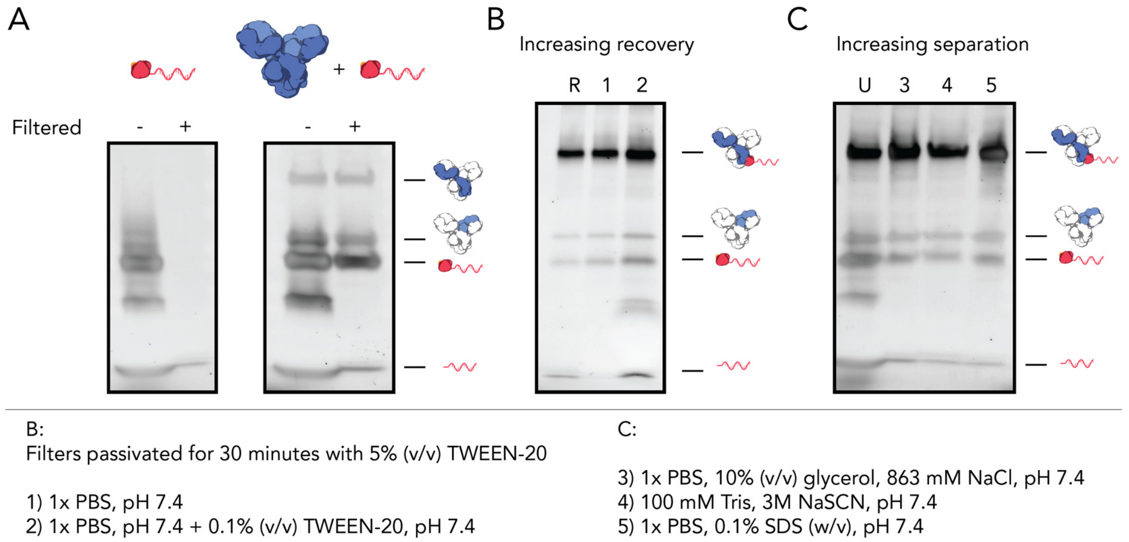


**Figure S8**. Optimization of pG-ODN-antibody purification using ultrafiltration. All SDS-PAGE gels were performed under reducing conditions and stained with SYBR Gold (A) pG-ODN removal using ultrafiltration in the absence (left) and presence of Cetuximab (right). Only in the absence of Cetuximab, pG-ODN was able to pass the 100 kDa molecular weight cut-off (MWCO) membrane. (B) pG-ODN removal using filters that were passivated with 5% TWEEN-20 for 30 minutes. When 0.1% (v/v) was included in the washing buffer, the recovery was increased to 68% compared to a reference filter that was not passivated (R). (C) pG-ODN removal using alternative (denaturing) washing buffers. In all cases most of the maleimide-ODN was removed successfully, however the separation between pG-ODN and pG-ODN-antibody constructs remained limited. Labels: U, unpurified sample.

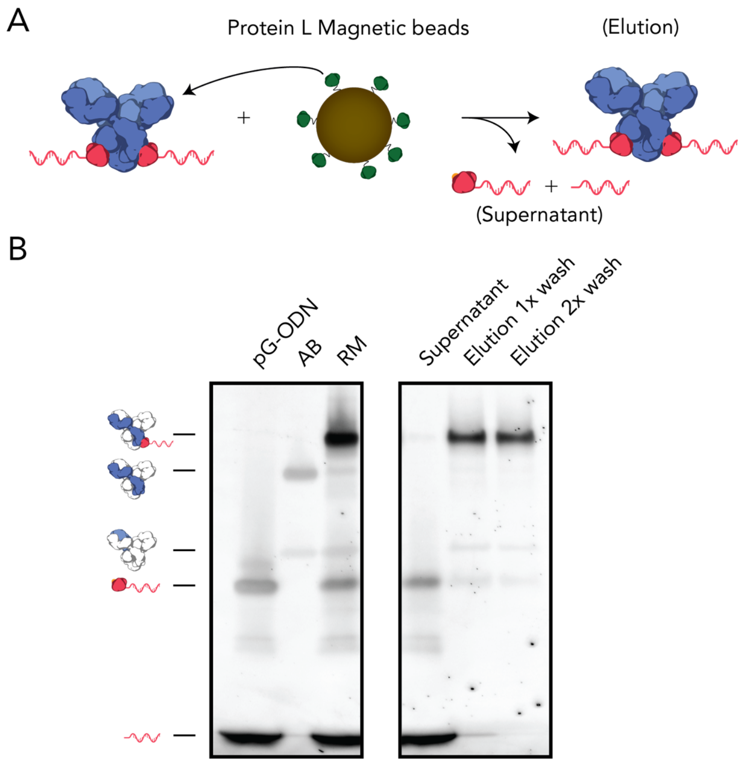


**Figure S9**. Purification of pG-ODN-Cetuximab using protein L magnetic beads. (A) Schematic overview of the purification process. Protein L binds to the light chain of Cetuximab and beads are recovered using a magnetic stand. (B) SDS-PAGE gel analysis under reducing conditions of different fractions of the purification. Both maleimide-ODN and pG-ODN remain in the supernatant while pG-ODN-antibody constructs are captured by the magnetic beads and recovered in the elution fraction. Labels: AB, Antibody (Cetuximab); RM, unpurified reaction mixture.

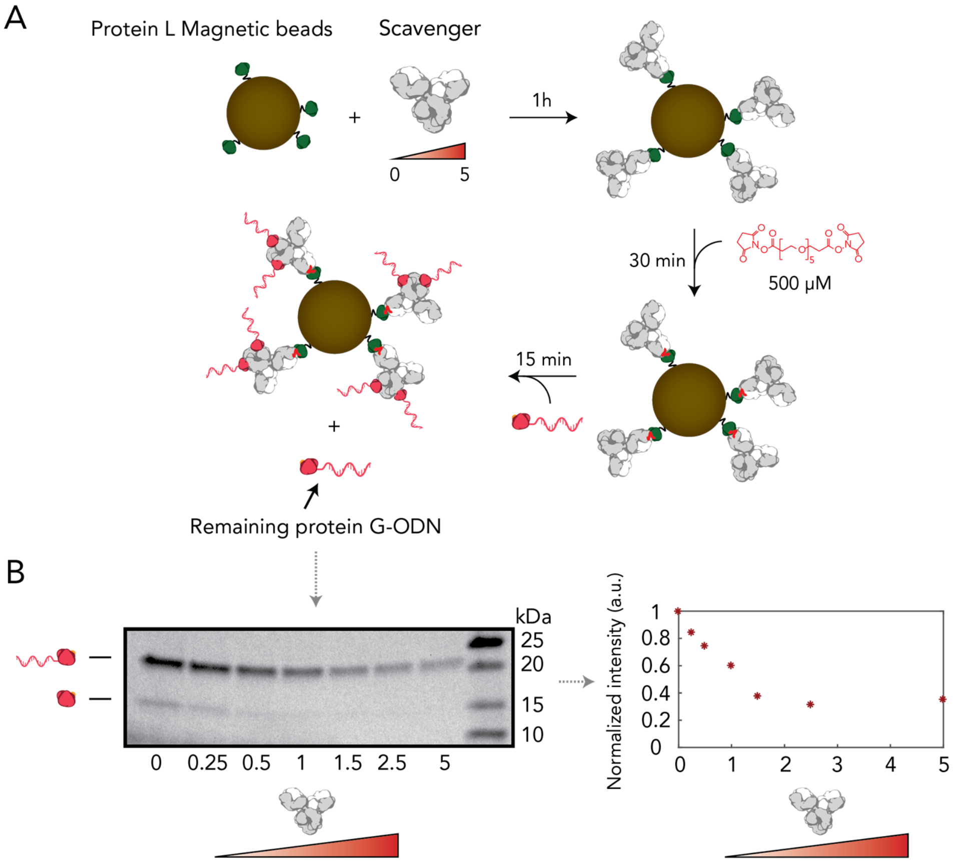


**Figure S10**. Optimization of the pG-ODN capturing capacity of the antibody-functionalized protein L magnetic beads as a function of the scavenger antibody. (A) Schematic overview of the synthesis of scavenging beads. Protein L magnetic beads were incubated with different equivalents of a scavenging antibody. In this case 1 equivalent was defined as the maximum capacity of a protein L bead (110 µg antibody/mg of bead). The antibody was covalently attached to the beads and the beads were incubated with pG-ODN. The beads were removed using a magnetic stand and the amount of remaining pG-ODN was quantified. (B) SDS-PAGE gel analysis under reducing conditions of the remaining amount of pG-ODN as a function of the amount of scavenging antibody. Gel band intensity shows that ≥ 2 equivalents of scavenger antibody are required to achieve maximum pG-ODN binding capacity.

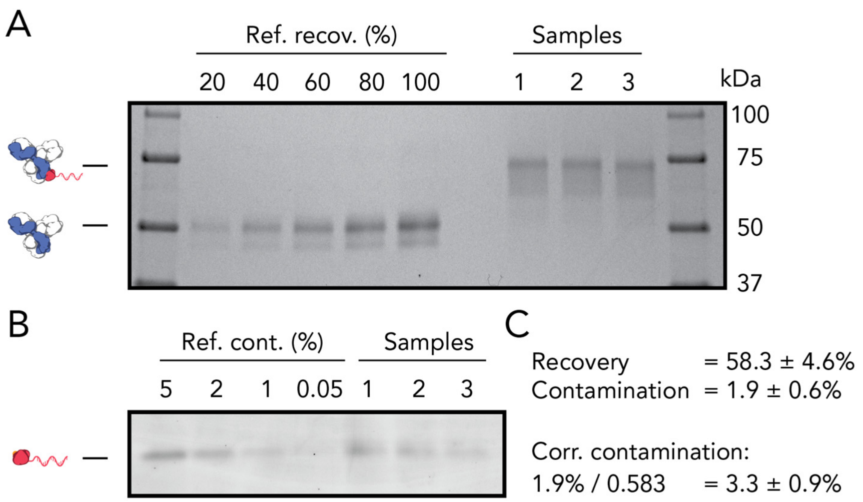


**Figure S11.** Recovery and purity of pG-ODN-antibody constructs purified using scavenging beads. (A) SDS-PAGE gel analysis under reducing conditions stained with Coomassie Blue. The gel band intensity corresponding to the recovered heavy chain of the pG-ODN-antibody was compared to a reference. (B) SDS-PAGE gel analysis under non-reducing conditions stained with SYBR Gold. The intensity of the band corresponding to pG-ODN after purification was compared to a reference. (C) Calculation of the recovery and contamination. The contamination was corrected based on the recovery of the pG-ODN-antibody construct.

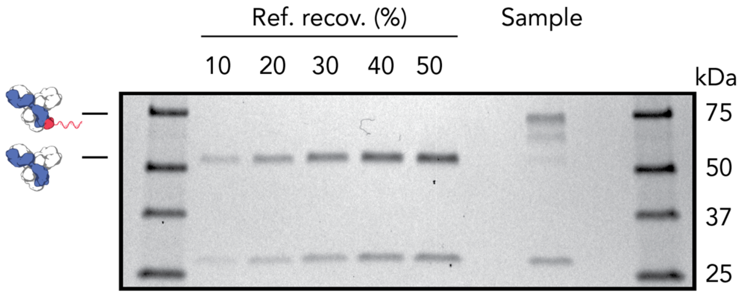


**Figure S12**. Recovery analysis of pG-ODN-Cetuximab using scavenging beads for purification. SDS-PAGE gel analysis under reducing conditions stained with Coomassie Blue. The gel band intensity corresponding to the recovered light chain of the pG-ODN-antibody was compared to a reference which showed a recovery of 30%

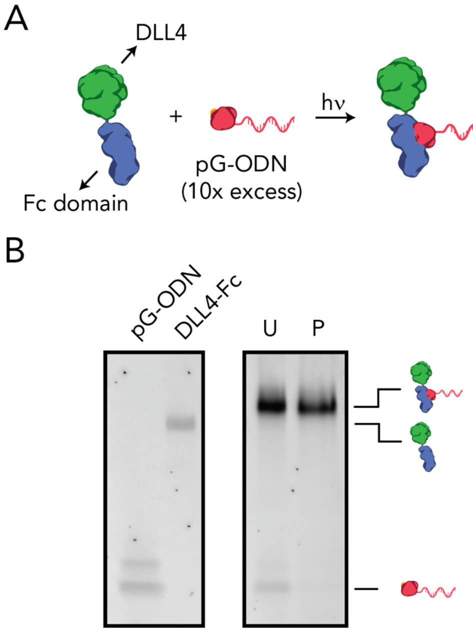


**Figure S13**. Labeling and purification of the Fc-fusion protein DLL4-Fc. (A) Schematic overview of the protein and labeling of DLL4-Fc using 10-fold molar excess of pG-ODN. (B) SDS-PAGE gel analysis under reducing conditions of unpurified (U) and purified (P) pG-ODN-DLL4-Fc using scavenging beads. The gel was stained with SYBR gold.

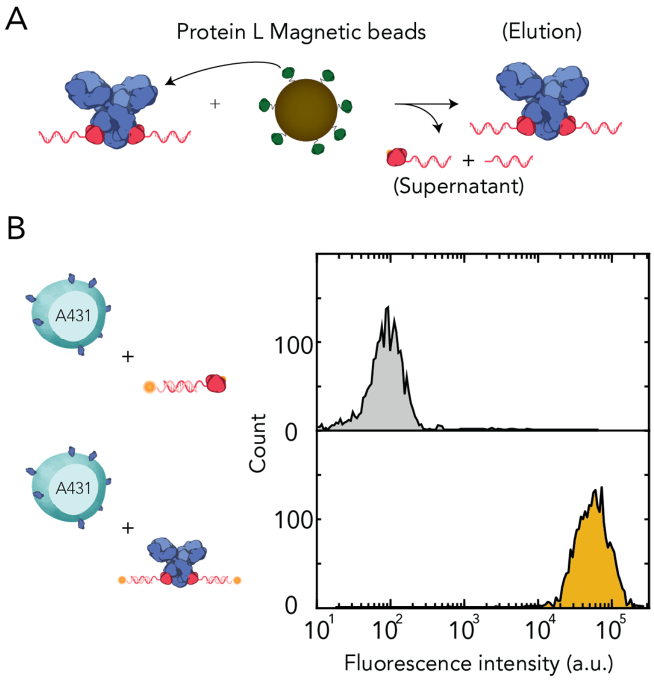


**Figure S14**. Antibody activity after direct purification of the pG-ODN-antibody construct using protein L functionalized magnetic beads. (A) Schematic overview of the purification process. Protein L binds to the light chain of Cetuximab and beads are recovered using a magnetic stand. (B) Flow cytometric analysis of EGFR-expressing A431 cells using 10 nM pG-ODN-functionalized Cetuximab hybridized to a CY5-labeled imager strand. Fluorescent intensity of pG-ODN-Cetuximab labeled A431 cells was compared to A431 cells incubated with only pG-ODN.

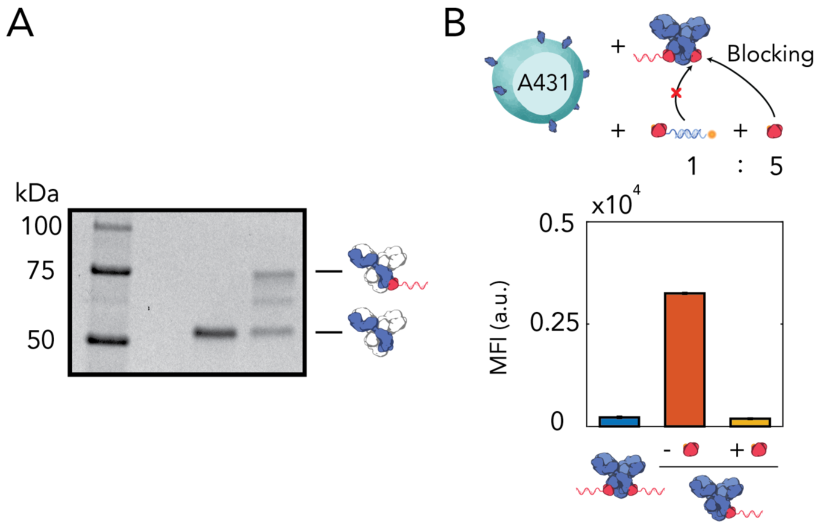


**Figure S15**. Blocking of free Fc sites using free pG during cellular labeling. (A) SDS-PAGE gel analysis under reducing conditions shows that ~50% of the Fc chains are covalently labeled with pG-ODN. (B) Partly labeled pG-ODN-Cetuximab was incubated with 20-fold molar excess of a competing pG-ODN sequence and 100-fold molar excess of pG to block free Fc sites. Flow cytometric analysis of EGFR-expressing A431 cells using 10 nM pG-ODN-functionalized Cetuximab shows that cross-contamination is observed when free Fc sites are not blocked with pG. Introduction of a 100-fold molar excess of pG shows the same median fluorescence intensity compared to quantitatively labeled pG-ODN-Cetuximab, indicating that successful blocking of Fc sites achieved. MFI represent the median fluorescence intensity and error bars represent SD (n=3).

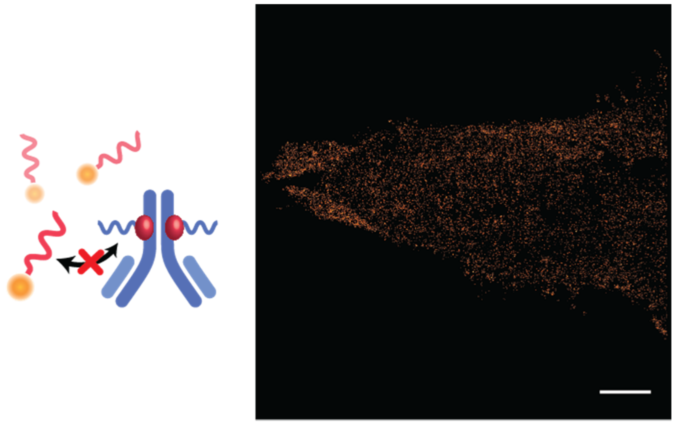

**Figure S16**. Aspecific interaction of imager strands in DNA-PAINT with cellular components. A431 carcinoma cells are labeled with a pG-ODN-Cetuximab construct containing a short (11 nt) docking strand (docking 4) and fixated to a glass slide. DNA-PAINT super-resolution image obtained using ATTO647N-functionalized imager strands which are not complementary to the docking strand (imager 5) (20,000 frames, 20-Hz frame rate). Scale bar, 5 µm.

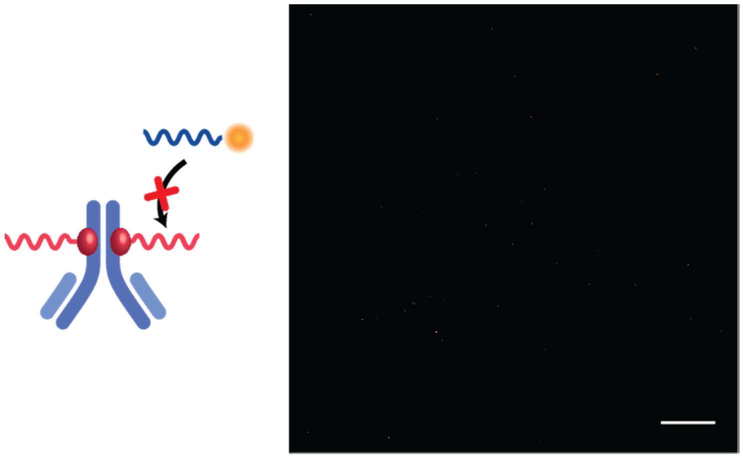

**Figure S17**. Aspecific interaction of imager strands in dSTORM with cellular components. A431 carcinoma cells are labeled with a pG-ODN-Cetuximab construct containing a long (20 nt) docking strand, not complementary to the imager (docking 5), and fixated to a glass slide. dSTORM super-resolution image obtained using CY5-functionalized imager strands (imager 1) (20,000 frames, 65.5-Hz frame rate). Scale bar, 5 µm.

**Supplementary materials and methods:**

**Antibodies/Fc-fusion proteins**

The following antibodies/Fc-fusion proteins were used to characterize the coupling of pG-ODN to antibodies from different host species: Cetuximab (anti-EGFR red.) (Erbitux, Merck), monoclonal anti-GFP mouse IgG2a (JL-8; catalog number: 632380, Clontech), Rabbit IgG Isotype Control (catalog number: 02-6102, Invitrogen) and DLL4-hIgG1-Fc (catalog number: 10171-H02H, Sino Biological). For the cellular labeling the following antibodies were used: monoclonal anti-CD45 mouse IgG2a (F10-89-4; catalog number: MA5-16669, Invitrogen) and monocolonal anti-CD31 mouse IgG2a (HEC7; catalog number: MA3100, Invitrogen)

**Cells and complementary culture medium**

A431 and Jurkat T cells were cultured in Roswell Park Memorial Institute (RPMI, Gibco) 1640 Medium fortified with 10% FBS and 1% penicillin and streptomycin. Human Umbilical Vein Endothelial Cells (HUVECs) were a kind gift from Dr. Cecilia Sahlgren. HUVECs were cultured in Endothelial Base Medium (EBM-2, Lonza) with additives of 2% FBS, 0.04% Hydrocortisone, 0.4% hFGF-B, 0.1% VEGF, 0.1% R3-IGF-1, 0.1% ascorbic acid, 0.1% hEGF, 0.1% GA-1000, and 0.1% heparin (Lonza) supplemented with 1% penicillin/streptomycin. All cells were incubated at 37 °C and 5% $CO_2$.

**SDS-PAGE**

For SDS-PAGE analysis 4-20% SDS-PAGE Mini-PROTEAN® TGX Precast gels (Bio-rad) were used (Novex® by Life Technologies). The running buffer consists of 25 mM Tris, 192 mM glycine, 0.1% SDS, pH 8.3. Samples were heated at 95 °C for 5 min in 1x SDS Sample Buffer (62.5 mM Tris, 10% Glycerol, 2.5% SDS (w/v), 0.01% Bromophenolblue, pH 6.8) before loading. When SDS-PAGE was performed under reducing conditions DTT to a final concentration of 50 mM was added. The gels were run at 150 V for either 40 minutes (analyzing pG-ODN coupling) or 60 minutes (analyzing pG-ODN-antibody labeling). To visualize DNA bands, the gel was stained in 50 mL 1x SYBR Gold (Thermo scientific) and to visualize protein bands Coomassie Brilliant Blue G-250 (Bio-Rad) was used.

**Buffer exchange/ODN removal using Amicon spin filters**

A spin-filter was pre-wetted with 500 µL of the buffer of interest and spun for 5 minutes at 14,000 xg at 4 °C. The remaining concentrate was removed after centrifugation. Before loading, the reaction mixture was diluted to a final volume of 500 µL in the buffer of interest. The sample was added to the filter and centrifuged for 5 minutes at 14,000 xg at 4 °C. This step was repeated for a total of three washing steps. Eventually, the desalted concentrate was recovered by inverting the filter and spinning for 6 minutes at 1,000 xg at 4 °C.

**Fast protein liquid chromatography (FPLC) with an anion-exchange column**

For FPLC an anion-exchange a HiTrap Q HP column (1 mL, GE Healthcare) was used. The column was equilibrated with equilibration buffer (50 mM Tris-HCl, pH 7.5). The conjugation reaction mixture was diluted five-fold in equilibration buffer and applied manually to the column. The column was washed with equilibration buffer supplemented with 100 mM NaCl and subsequently a salt gradient was applied with a start and end concentration of 100 and 500 mM NaCl, respectively, and a flow rate of 1 mL/min. Elution fractions of 0.5 mL were collected and analyzed by measuring on-line absorption at 280 nm.

**Quadrupole time-of-flight mass spectrometry (Q-Tof)**

For Q-Tof an aliquot of pG was buffer exchanged using Amicon 3 kDa MWCO centrifugal filters (Merck Millipore) to ultrapure $H_2O$ to a final concentration of 1 mg/mL. A sample of 0.1 µL pG was injected to an Agilent Polaris C18A RP column with a flow of 0.3 mL/min and a 15-60% acetonitrile gradient containing 0.1% formic acid. Mass spectra were measured on a Xevo G2 QTof mass spectrometer (Waters) in positive mode and deconvoluted with MaxEnt Deconvolution software.

**Size exclusion chromatography (SEC)**

Size exclusion chromatography was performed on an Agilent 1260 Infinity II Bio-inert LC System using a Bio-SEC-5 column (5 µm, 300 Å, 7.8*300 mm, Agilent). The column was equilibrated with 5 column volumes of 1x PBS, pH 7.4 using a flow rate of 1 mL/min. The column was coupled to a 280 nm UV spectrophotometer and a fraction collector. 95 µL sample was injected and elution fractions of 0.2 mL were collected and analyzed on SDS-PAGE under reducing conditions

**Direct purification using protein L beads**

This procedure is optimized to purify 50 µL of 4 µM antibody-pG-ODN from 40 µM uncoupled pG-ODN using 25 µL of protein L beads. In a typical reaction 25 µL of Pierce™ protein L magnetic beads was added to 75 µL washing buffer (1x PBS, 0.05% TWEEN-20, pH 7.4). The beads were washed twice in 200 µL washing buffer after which the magnetic beads were incubated with 50 µL of the reaction mixture diluted to a final volume of 200 µL in washing buffer. Reaction was rotated at for 1h at 4 °C. Subsequently, the magnetic beads were collected with a magnetic stand and the supernatant was discarded. The protein L beads were washed twice using 200 µL washing buffer. To collect the antibody-pG-ODN, beads were incubated for 5 minutes in elution buffer (100 mM Tris-HCl, 1 M NaSCN, pH 7.5). The elution fraction was desalted using a Zeba™ spin desalting column, 7000 MWCO, 0.5 mL (Thermo scientific).

**Blocking of free Fc sites**

Cetuximab was incubated for 1h under UV-illumination using 3-fold molar excess of pG-ODN to achieve 50% labeling of Fc domains. 20-fold molar excess of a competing pG-ODN was added to the partly labeled pG-ODN-Cetuximab in the presence and absence of 100-fold molar excess pG. The reaction mixture was shaken at 600 RPM for 1h at 20 °C. Subsequently, 12.5 µL A431 cells ($3.5*10^6$ cells/mL) were incubated with the reaction mixture in a final reaction volume of 250 µL and a final concentration of 10 nM pG-ODN-Cetuximab. Incubation was performed at 400 RPM for 30 minutes at 20 °C. After incubation the cells were pelleted and redissolved in labeling buffer (1x PBS, 0.1% (w/v) BSA, pH 7.4) containing 1 µM pG and 100 nM of a CY5-labeled ODN complementary to the competing pG-ODN sequence. The cells were incubated and centrifuged as described above and analyzed using flow cytometry.

**Flow cytometry**

Flow cytometry was performed on a FACS Aria III (BD Biosciences) equipped with a 70 µm nozzle. Single cell events were gates based on the linear relation between forward scatter height (FSC-H) and forward scatter area (FSC-A). For each sample 2,000 gated events were collected and analyzed using custom written Matlab scripts.

**Table 1: DNA sequences**

All DNA oligonucleotides were purchased from Integrated DNA Technologies. Amino-functionalized ODNs were obtained desalted and dissolved in DNas/Rnase-free water at a concentration of 1 mM and fluorescently labeled ODNs were HPLC purified and dissolved at a concentration of 250 µM

**SDS-PAGE analysis pG-ODN/antibody coupling**

| Name | Sequence (5' to 3') | Length (nt) |
|---|---|---|
| Docking 1 (Cetuximab) | CCC TAG AGT GAG TCG TAT GA/3AmMO/ | 20 |

**Flow cytometry**

| Name | Sequence (5' to 3') | Length (nt) |
|---|---|---|
| Imager 1 (Cetuximab) | TCA TAC GAC TCA CTC TAG GGT T/3Cy5Sp/ | 22 |
| Docking 2 (aCD45) | /5AmMC6/AC TGA CTG ACT GAC TGA CTG | 20 |
| Imager 2 (aCD45) | /5Cy5/CA GTC AGT CAG TCA GTC AGT | 20 |
| Docking 3 (aCD31) | GTC CAT GCT CAG GAT TGC GA /3AmMO/ | 20 |
| Imager 3 (aCD31) | TCG CAA TCC TGA GCA TGG ACT T /3Cy5Sp/ | 22 |

**DNA-PAINT & dSTORM**

| Name | Sequence (5' to 3') | Length (nt) |
|---|---|---|
| Docking 4 (DNA-PAINT) | /5AmMC6/TTA TAC ATC TA | 11 |
| Imager 4 (DNA-PAINT, c.[1]) | CTA GAT GTA T/3ATTO647NN/ | 10 |
| Imager 5 (DNA-PAINT, n.c. [2]) | TAT GTA GAT C/3ATTO647NN/ | 10 |
| Docking 5 (dSTORM, n.c.[2]) | TTA TAC ATC TAG TCG TGT GA/3AmMO/ | 20 |

[1] complementary
[2] non-complementary

For dSTORM imaging docking 1 and imager 1 were used to generate figure 5.

**Table 2: Protein G sequences**

The single-letter amino acid code is shown in uppercase above the corresponding DNA sequence. The cysteine is shown in purple, the *strep*-tag in orange, pG in blue, the hexahistidine tag in green and the amber stop codon, coding for the non-natural amino acid *p*-Bpa, is indicated in red.

**pG-C2:**

```
      M  C  W  S  H  P  Q  F  E  K  G  T  M  T  F  K  L  I  I  N
1     ATGTGCTGGTCCCATCCGCAGTTCGAGAAAGGTACCATGACATTTAAACTGATAATCAAC

      G  K  T  L  K  G  E  I  T  I  E  A  V  D  A  *  E  A  E  K
61    GGCAAAACCTTAAAAGGGGAGATCACAATTGAGGCAGTCGATGCCTAGGAAGCCGAGAAA

      I  F  K  Q  Y  A  N  D  Y  G  I  D  G  E  W  T  Y  D  D  A
121   ATCTTTAAACAATATGCTAATGATTATGGTATTGACGGAGAATGGACGTATGACGATGCG

      T  K  T  F  T  V  T  E  E  F  T  S  G  G  S  G  D  D  H  H
181   ACAAAAACTTTCACCGTAACTGAGGAATTCACTAGTGGTGGAAGTGGGGACGATCATCAT

      H  H  H  H  *
241   CATCATCATCATTAA
```

**pG-C3:**

```
      M  S  C  W  S  H  P  Q  F  E  K  G  T  M  T  F  K  L  I  I
1     ATGAGTTGCTGGTCCCATCCGCAGTTCGAGAAAGGTACCATGACATTTAAACTGATAATC

      N  G  K  T  L  K  G  E  I  T  I  E  A  V  D  A  *  E  A  E
61    AACGGCAAAACCTTAAAAGGGGAGATCACAATTGAGGCAGTCGATGCCTAGGAAGCCGAG

      K  I  F  K  Q  Y  A  N  D  Y  G  I  D  G  E  W  T  Y  D  D
121   AAAATCTTTAAACAATATGCTAATGATTATGGTATTGACGGAGAATGGACGTATGACGAT

      A  T  K  T  F  T  V  T  E  E  F  T  S  G  G  S  G  D  D  H
181   GCGACAAAAACTTTCACCGTAACTGAGGAATTCACTAGTGGTGGAAGTGGGGACGATCAT

      H  H  H  H  H  *
241   CATCATCATCATCATTAA
```
